# Remote neuronal activity drives glioma infiltration via Sema4f

**DOI:** 10.1101/2023.03.15.532832

**Authors:** Emmet Huang-Hobbs, Yi-Ting Cheng, Yeunjung Ko, Estefania Luna-Figueroa, Brittney Lozzi, Kathryn R Taylor, Malcolm McDonald, Peihao He, Hsiao-Chi Chen, Yuhui Yang, Ehson Maleki, Zhung-Fu Lee, Sanjana Murali, Michael Williamson, Dongjoo Choi, Rachel Curry, James Bayley, Junsung Woo, Ali Jalali, Michelle Monje, Jeffrey L Noebels, Akdes Serin Harmanci, Ganesh Rao, Benjamin Deneen

## Abstract

The tumor microenvironment (TME) plays an essential role in malignancy and neurons have emerged as a key component of the TME that promotes tumorigenesis across a host of cancers. Recent studies on glioblastoma (GBM) highlight bi-directional signaling between tumors and neurons that propagates a vicious cycle of proliferation, synaptic integration, and brain hyperactivity; however, the identity of neuronal subtypes and tumor subpopulations driving this phenomenon are incompletely understood. Here we show that callosal projection neurons located in the hemisphere contralateral to primary GBM tumors promote progression and widespread infiltration. Using this platform to examine GBM infiltration, we identified an activity dependent infiltrating population present at the leading edge of mouse and human tumors that is enriched for axon guidance genes. High-throughput, *in vivo* screening of these genes identified Sema4F as a key regulator of tumorigenesis and activity-dependent infiltration. Furthermore, Sema4F promotes the activity-dependent infiltrating population and propagates bi-directional signaling with neurons by remodeling tumor adjacent synapses towards brain network hyperactivity. Collectively, our studies demonstrate that subsets of neurons in locations remote to primary GBM promote malignant progression, while revealing new mechanisms of tumor infiltration that are regulated by neuronal activity.

---

Glioblastoma (GBM) is the most aggressive and lethal form of brain tumor, featuring high rates of proliferation and infiltration into surrounding brain tissue^1–4^. Despite treatment, recurrence is inevitable and tends to occur outside surgical margins or in locations remote to the primary tumor^5–7^, highlighting the central role that tumor infiltration plays in this malicious disease. GBM infiltration in the brain generally occurs along organized anatomical structures such as blood vessels and white matter tracts, which contain neuronal axons and suggests the involvement of neuronal populations^8–11^. Previous studies established correlations between the presence of GBM and heightened neuronal activity in surrounding brain regions^12–16^. Moreover, it has been shown that increased neuronal activity can promote optic nerve glioma progression and the growth of both pediatric and adult forms of high-grade glioma through mechanisms involving activity-regulated paracrine factors and neuron-to-glioma synaptic signaling^16–19^. This raises the possibility that neuronal activity itself can promote tumor infiltration, a concept supported by the recent discovery that direct synaptic signaling between neurons and glioma cells can promote invasion^20,21^. Whether neuronal activity promotes circuit-specific patterns of glioma infiltration through paracrine signaling is unknown, and the underlying molecular mechanisms driving GBM infiltration remain obscure. Furthermore, the brain contains a plethora of neuronal subtypes, and which subtypes of neurons serve as the substrate for driving GBM progression is also incompletely understood.

## Results

### Contralateral neuronal stimulation promotes glioma progression

Neuronal activity promotes glioma proliferation, however whether activity promotes transformation of low-grade glioma (LGG) to high-grade glioma (HGG) remains an open question^17,22^. Furthermore, these prior studies stimulated neurons in close proximity to xenografted tumors, raising the question of whether long-range neuronal projections from brain regions remote to the primary tumor also contribute to tumorigenesis. To determine whether remote stimulation of neurons promotes LGG to HGG transformation, we used the native RCAS/Ntva system that is driven by overexpression of PDGFB and generates LGG^23,24^. After initiating tumors in the cortex at P1, we injected the contralateral cortex with AAV2/9 Syn1-hM3Dq-mCherry at P5 (**Fig.1a**). To stimulate contralateral neurons, we treated mice with saline or 5 mg/Kg of clozapine N-oxide (CNO) two times a day, for two months, starting at P20. To confirm that CNO treatment activates neurons contralateral to the tumor, we performed slice recordings and found increased activity upon CNO treatment (**Extended Data Figure 1a-b**). Strikingly, mice treated with CNO exhibited a drastic decrease in median overall survival compared to the saline group (51days CNO v. 95days saline) (**Fig.1b**). These changes in survival are complemented by increased Ki67 expression in the CNO group and coupled with hallmark pathological features of HGG, including microvascular proliferation and necrosis (**Fig.1c-red arrows**). These observations indicate that stimulation of neuronal activity in regions remote to primary LGG can promote progression to HGG.

**Figure 1.**
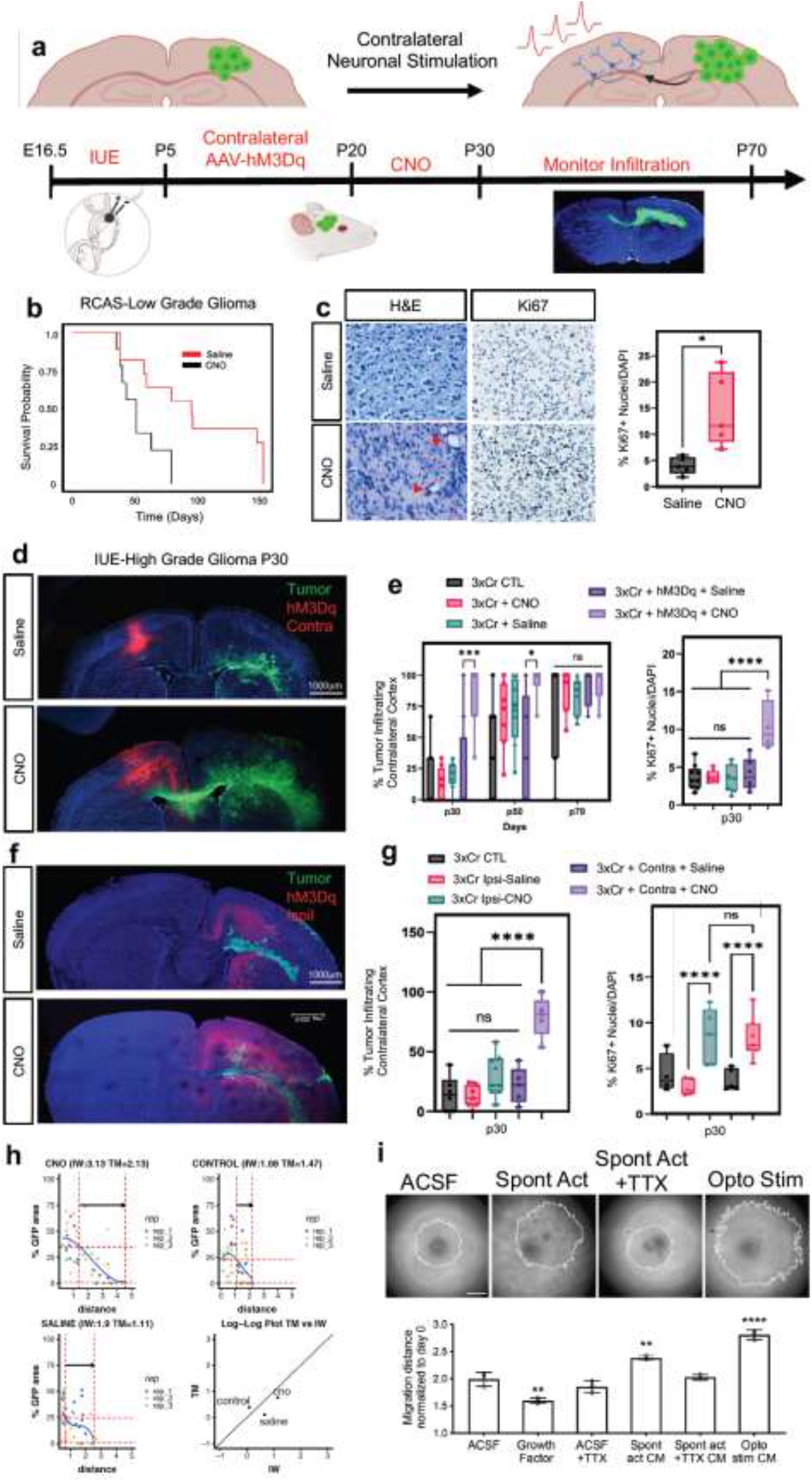
Remote neuronal stimulation accelerates glioma progression. **a.** Schematic of DREADD-based activation of neurons contralateral to tumor in both RCAS-Ntva and IUE models. **b.** Kaplan-Meier survival analysis of RCAS-Ntva tumors treated with saline (median_Saline_= 95 days, n = 11) or CNO (median_CNO_ = 51 days, n = 9) showing significantly faster morbidity in CNO treated RCAS tumors (Log-rank (Mantel-Cox) test, Chisq = 6.456, df = 1, p-value = 0.0111, CNO/Saline HR_log-rank_= 2.768, 95% CI = 0.9770 to 7.945). **c.** H&E staining of RCAS-Ntva tumors samples revealed high grade characteristics in CNO treated tumor groups (red arrows). Ki67 staining proliferation in CNO treated mice versus saline treated mice. Quantification is derived from n=5 mice from CNO (mean = 14.51%, SD = 7.076%) and Saline (mean = 4.017%, SD = 2.179%) groups and determined by Welch’s unpaired t-test (p-value = 0.0276, t = 3.228, df = 4.441). **d.** Representative images from IUE-HGG tumors at P30 demonstrating infiltration; green is tumor, red is AAV-DREADD virus. **e.** Quantification of infiltration and Ki67 expression across the P30-P70 timecourse. Infiltration was quantified based on the presence of tumor cells in contralateral cortex and analyzed via two-way ANOVA; data derived from p30 CTL n = 8, p50 CTL n = 7, p70CTL n = 7, p30+CNO n = 8, p50+CNO n = 7, p70+CNO n = 7, p30+Saline n = 8, p50+Saline n = 7, p70+Saline n = 7, p30+AAV+Saline n = 8, p50+AAV+Saline n = 5, p30+AAV+Saline n = 4, p30+AAV+CNO n = 8, p50+AAV+CNO n = 6, p30+AAV+CNO n = 5 samples. Ki67 staining was performed at the p30 time point, from CTL n = 9, CNOonly n = 8, Salineonly n = 7, AAV+Saline n = 5, AAV+CNO n = 4 samples. **f.** Representative images from IUE-HGG tumors at P30 demonstrating the extent of infiltration after activation of neurons in the cortex ipsilateral to the tumor (CNO) and saline treated controls; green is tumor, red is AAV-DREADD virus. **g.** Quantification of tumor infiltration and Ki67 expression at P30 (3xCr CTL n = 5 mean = 4.329%, 3xCr+IpsilAAV+Saline n=5, mean = 2.975%, 3xCr+IpsilAAV+CNO n=5, mean = 8.144%). Significant difference was found in Ki67+ nuclei in ipsilateral CNO stimulated tumors vs saline control (p-value <0.0001) and vs 3xCr only controls (p-value = 0.0020); Infiltration was quantified based on the presence of tumor cells in contralateral cortex and analyzed via one-way ANOVA (3xCr CTL n = 6, 3xCr+IpsilAAV+Saline n=7, 3xCr+IpsilAAV+CNO n=9) with CNO stimulated brains showing no statistical difference to Saline treated (p-value = 0.0649) or control tumors (p-value = 0.1504). Direct comparison between ipsilateral-CNO and contralateral-CNO groups revealed a statistically significant difference (p-value <0.0001). **h.** Mathematical modeling of glioma infiltration as a function of tumor mass. Blue line is the smoothed data points using piecewise-cubic splines; red horizontal dashed lines are the 0.8 *p*_max_ and 0.02 *p*_max_ glioma cell density of the maximum smoothed cellular density (*p*_max_). Red vertical lines are the intersecting distance points of the red horizontal lines with smoothed blue line, which is used in calculating infiltrating width (IW). Black arrow shows the IW. Log-log plot shows the dependence of IW and tumor mass (TM). Analysis was performed at the p30 timepoint on CTL n = 3, Saline n = 3, CNO n = 3; samples from individual biological replicates are color coded. **i.** Glioma 3D spheroid migration assay, measuring glioma infiltration after treatment with growth factor media, conditioned media (CM) from spontaneously active cortical explants, spontaneously active cortical explants silenced with TTX (10μm) or optogenetically stimulated cortical explants (channelrhodopsin-2 (ChR2)-expressing deep layer cortical projection neurons), in comparison to ACSF control. Scale bar is 500μm. \**P* < 0.05, \*\**P* < 0.01, \*\*\**P* < 0.001, **** *P* < 0.0001, log-rank(**b**), unpaired Welch’s t-test (**c**), two-way analysis of variance (ANOVA) (**e**), one-way analysis of variance (ANOVA) (**c, e**).

Infiltration throughout the brain is another key facet of HGG progression, which we examined using our native CRISPR/Cas9-based in utero electroporation (IUE) model of HGG. Using IUE-based approaches we introduced gRNAs to *NF1*, *PTEN*, and *p53* into a single cortical ventricle at e16.5, which initiates glioma tumorigenesis in a single cortical hemisphere and eventually infiltrates across the corpus callosum to the contralateral hemisphere (**Fig. 1a and Extended Data Figure 1c-d**). To examine whether stimulation of neurons from brain regions remote to the primary tumor promotes infiltration, we injected the contralateral cortex with AAV2/9 Syn1-hM3Dq-mCherry at P5 (**Fig.1a)** and treated with CNO (or saline controls) starting at P20. Using migration across the corpus callosum, into the contralateral cortex as an index of glioma infiltration, we found that CNO treated tumors exhibited a dramatic increase in infiltration as early as P30 (i.e., 10 days post-CNO) and was also observed at P50 (**Fig.1d-e**). This acceleration of glioma infiltration was complemented by an increase in Ki67 expression in the CNO group (**Fig.1e and Extended Data Figure 1e**); critically CNO-only controls (without hM3Dq) did not demonstrate any effects on either tumor proliferation or infiltration (**Fig.1e and Extended Data Figure 1f-h**). To ascertain whether these effects on infiltration are specific to stimulation of contralateral neurons, we generated tumors and activated neurons in the ipsilateral cortex, finding an increase in proliferation, but no significant changes in infiltration at P30 when compared to saline controls. (**Fig.1f-g and Extended Data Figure 2a**). Moreover, direct comparison between contralateral and ipsilateral stimulation groups revealed a significant increase in infiltration after contralateral stimulation (**Fig.1g**). Together, these findings suggest that neuronal activity in the hemisphere contralateral to the primary tumor specifically promotes precocious tumor infiltration across cortical hemispheres.

To confirm that stimulation of contralateral neurons promotes glioma infiltration, we employed mathematical modeling^25,48^ finding that tumor infiltration width (IW) is increased relative to tumor mass (TM) in the CNO treated group compared to the saline group and non-treated control tumors at P30 (**Fig.1h**). To independently validate that neuronal activity-regulated paracrine factors promotes glioma infiltration, we used patient-derived glioma cell cultures in conjunction with an established three-dimensional spheroid culture system to measure infiltration ^26,27^. These cultures were treated with conditioned media (CM) from cortical explants with spontaneously active neurons, optogentically stimulated neurons, unconditioned control media (artificial cerebrospinal fluid, ACSF), or CM from cortical explants treated with TTX to silence neuronal activity. These studies revealed that treatment of glioma spheroids with CM from cortical explants containing active neurons promoted glioma infiltration (**Fig.1i**). Collectively, these data indicate that neuronal activity promotes tumor infiltration through secreted factors and that neurons contralateral to the primary tumor specifically drive this phenomenon during the early stages of tumor progression.

### Callosal projection neurons promote glioma progression

The preceding observations raise the question of which neuronal populations in the contralateral cortex are driving glioma infiltration and progression. Given that the axons of callosal projection neurons (CPN’s) cross cortical hemispheres, we reasoned that this population in the contralateral cortex is contributing to activity dependent infiltration^28^. CPN axons cross the cortical hemisphere along the corpus callosum, a white matter tract that also serves as a major route of glioma infiltration. Therefore, to test whether CPN’s are necessary for driving activity dependent tumor infiltration in our system, we severed the corpus callosum in the context of stimulation of contralateral neurons. Using our established paradigm (**Fig.1a)**, we severed the corpus collosum at P10 and initiated CNO and saline treatments at P20 (along with non-severed controls), followed by harvesting tumor bearing brains at P30. Analysis of these tumors revealed that severing the corpus callosum abolished the activity-dependent acceleration of infiltration that was observed with the intact control (**Fig.2a-b**). Moreover, we observed that activity-dependent increases in Ki67 expression were also lost after severing the corpus callosum (**Fig.2b and Extended Data Figure 2b).** These results indicate that an intact corpus callosum is necessary for contralateral neurons to promote glioma progression, implicating CPNs in this phenomenon.

**Figure 2.**
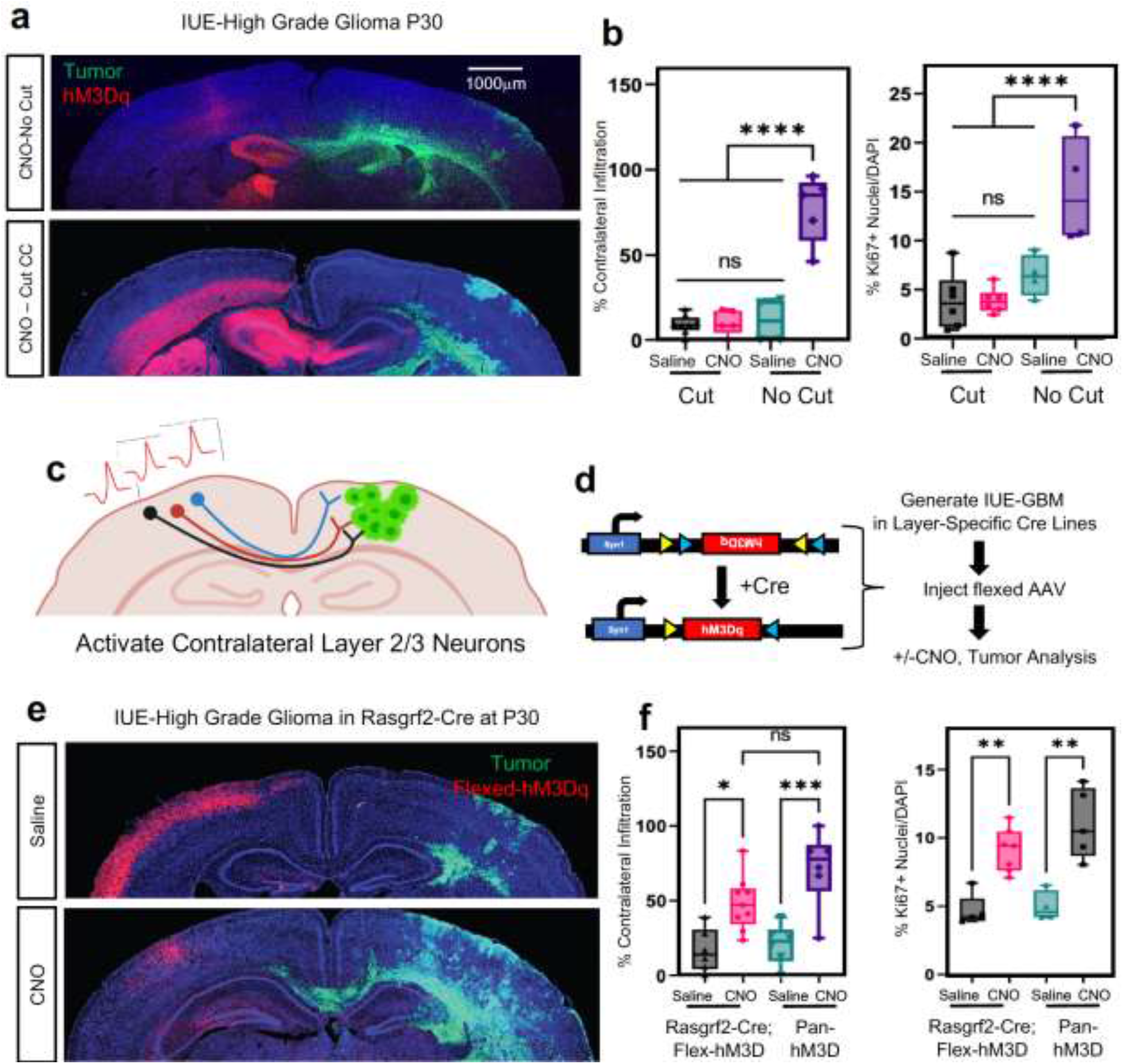
Callosal projection neurons promote glioma infiltration. **a.** Representative images from IUE-HGG tumors at P30 demonstrating the extent of infiltration with a severed corpus callosum (cut) or control (no cut); green is tumor, red is AAV-DREADD virus. **b.** Quantification of tumor infiltration and Ki67 expression at P30, data derived from CTL n = 4, AAV+CNO n = 5, AAV+Saline n = 4, CCcut+AAV+CNO n = 6, CCcut+AAV+Saline n = 6, Infiltration was quantified based on the presence of tumor cells in contralateral cortex and analyzed via ordinary one-way analysis of variance (ANOVA); CTL n=8, CC-Cut+AAV+CNO n=5, CC-cut+AAV+Saline n=8 (≥12 coronal sections assessed per brain). **c.** Schematic of callosal projection neuron activation experiment. **d**. Schematic of combined Rasgrf2-dCre mouse line with Cre-inducible DIO-hM3D-2a-mCherry DREADD to selectively activate layer2/3 neurons in contralateral hemisphere. **e.** Representative images from IUE-HGG tumors, injected with AAV-DIO-hM3D-2a-mCherry in Rasgrf2-Cre mice. Mice were harvested at P30 and the extent of tumor infiltration was evaluated **f.** Quantification of tumor infiltration and Ki67 expression at P30; Infiltration was quantified based on the presence of tumor cells in contralateral cortex and analyzed via ordinary one-way analysis of variance (ANOVA); Rasgrf2+AAV+CNO n=9, Rasgrf2+AAV+Saline n=6, 3xCr+AAV+CNO n=6, 3xCr+AAV+Saline n=6 (≥12 coronal sections assessed per brain). Proliferation samples were analuzed via one was analysis of variance; Rasgrf2+AAV+CNO n=7, Rasgrf2+AAV+Saline n=5, 3xCr+AAV+CNO n=5, 3xCr+AAV+Saline n=4. \**P* < 0.05, \*\**P* < 0.01, \*\*\**P* < 0.001, **** *P* < 0.0001, one-way analysis of variance (ANOVA) (**b, f**).

To examine whether CPNs are sufficient to accelerate tumor progression we utilized the Rasgrf2-dCre line, which marks layers 2/3 of the cortex and from which ~80% of CPN’s are derived (**Fig.2c**)^28,29^. To achieve selective activation of Rasgrf2-dCre expressing neurons, we utilized a double-floxed inverse orf (DIO) construct (pAAV-Syn1-DIO-hM3D-2A-mCherry), while inducing dCre with Trimethorpim, at 100ng/g body weight (**Fig.2d**). Induction of dCre and activity of *Rasgrf2-Cre; ROSA-floxed-tdTomato* in cortical layer 2/3 was confirmed (**Extended Data Figure 3a**), enabling us to use this mouse line in our IUE-based, glioma-activity paradigm (**Fig.1a**). Here, we injected the AAV-DIO virus in the contralateral cortex at P5, followed by dCre induction at P15, which enabled expression of hM3Dq in layer 2/3 neurons. After these manipulations, we treated mice with saline or CNO at P20 and harvested tumor bearing brains at P30. Strikingly, selective stimulation of Rasgrf2-Cre expressing neurons with CNO in the contralateral hemisphere promoted both tumor infiltration and Ki67 expression when compared to saline controls (**Fig.2e-f and Extended Data Figure 2c)** and at levels comparable to pan-neuronal activation controls. As a control for the specificity of this manipulation we activated inhibitory neurons, which are distinct from CPNs, in the contralateral hemisphere using AAV2/9 Dlx5/6-hM3Dq-mCherry in our paradigm. These studies revealed no changes in tumor cell infiltration or proliferation at P30, further supporting the specificity of Layer2/3 neurons from the contralateral hemisphere (**Extended Data Figure 3b-c**). Collectively, these data indicate that CPN’s play a critical role in driving tumor progression, while highlighting the contributions of neurons in remote brain regions to glioma progression.

### Identification of activity-dependent, infiltrating glioma populations

Infiltrating glioma cells play a central role in progression and eventual recurrence. Our activity-driven paradigm of glioma progression offers a venue in which to examine the cellular and molecular properties of these critical, yet poorly defined populations. To achieve this, we performed single-cell RNA-sequencing on P50 glioma tumors generated in the presence of contralateral stimulation (and saline controls) and using GFP as a marker of tumor cells we were able to distinguish host microenvironmental populations from tumor populations (**Fig.3a**). This analysis revealed widespread changes in the immune microenvironment in the presence of increased neuronal activity (**Fig. 3a and Extended Data Figure 3d-e and Extended Tables 1-2**), coupled with changes in the cellular constituency of the tumor. Focusing on the GFP+ tumor populations, we performed additional analysis and identified several prospective subpopulations that are enriched in the CNO, stimulated tumors (**Fig.3b**). We performed Gene Ontology (GO) analysis on the most enriched subpopulation in this CNO-enriched clusters (**Fig.3b-red arrow**) and identified a host of unique GO terms, including genes associated with glutamatergic synapses and axon guidance (**Fig.3b and Extended Table 3**). Next, we sought to localize this activity-dependent subpopulation within the tumor, hypothesizing that it likely resides at the leading edge in the contralateral hemisphere. Using spatial transcriptomics of P50 activity-driven glioma, we localized the gene signatures associated with the activity-dependent cluster in infiltrating tumor cells in the cortex contralateral to the primary tumor (**Fig.3c-d**). Furthermore, the cells at the leading edge were also enriched for genes associated with axon guidance (**Extended Data Figure 3f and Extended Table 4**). These observations in mouse models led us to examine whether this infiltrating population is also present in the leading edge of human GBM. Therefore, we cross-correlated the gene signature associated with the infiltrating mouse population with the IVY-GAP database, which has transcriptomic data for distinct anatomical structures of GBM, including the leading edge^30^. This analysis revealed a similar and highly specific enrichment of this infiltrating signature at the leading edge of human GBM (**Fig.3e).** Together these observations indicate that neuronal stimulation drives the generation of infiltrating populations, and these populations correspond to the leading edge of GBM tumors.

**Figure 3.**
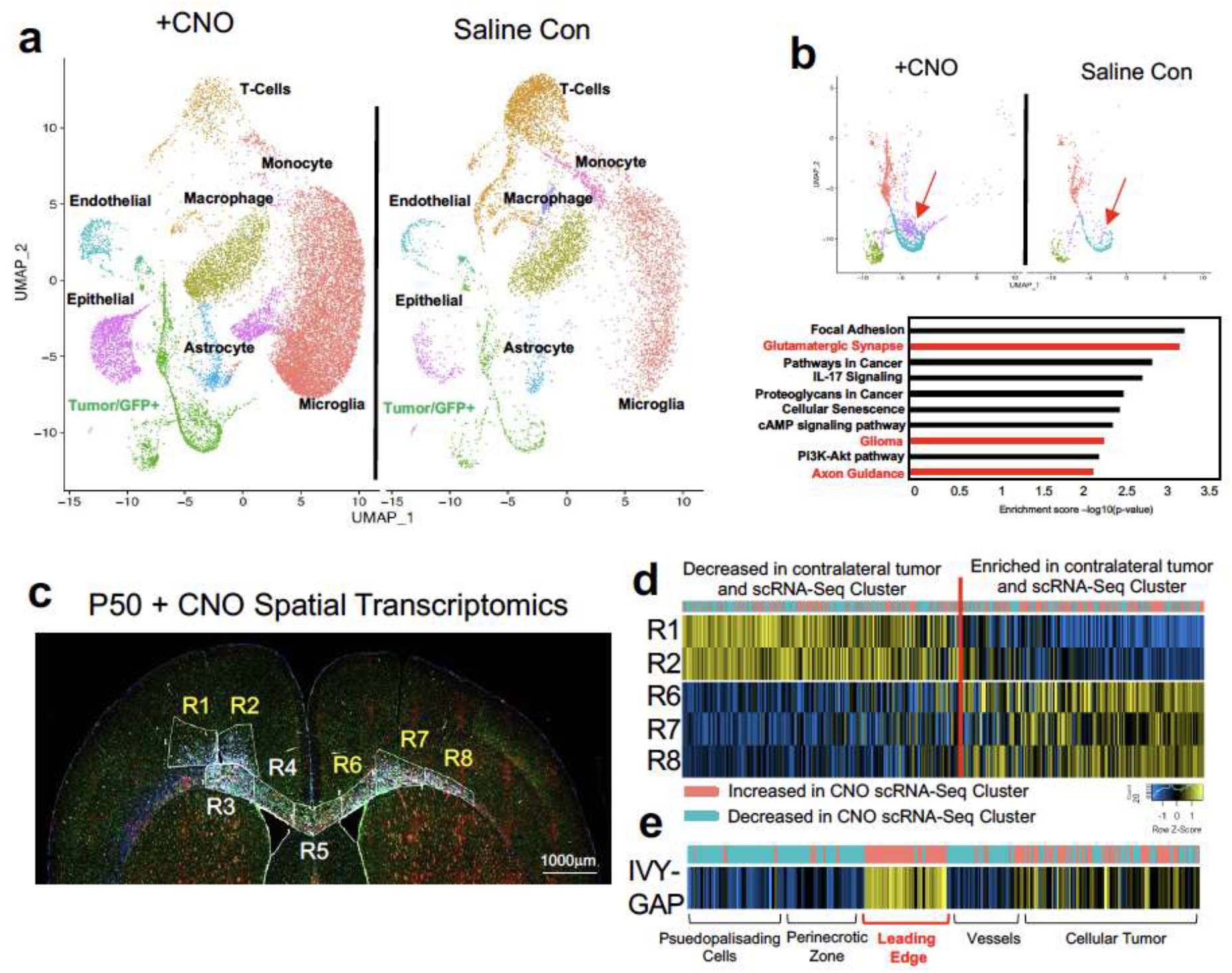
Identification of activity-dependent infiltrating glioma population. **a**. Single Cell RNA-Seq DimPlots of P50 IUE-HGG from CNO and saline controls. Cell types were mapped in SingleR with celldex as expression profile reference. **b.** Sub-clustering analysis of GFP+ tumor cells shown in **a**. red arrow indicates the cluster of interest we chose to investigate. GO-term analysis on this prospective cell cluster was performed using EnrichR and the KEGG 2021 Human dataset. **c.** Spatial transcriptomics on P50 IUE-HGG from the CNO group were performed with GeoMx Nanostring Digital Spatial Profiling. Regions were selected based on the expression of elevated GFAP and vimentin in the section and the presence of GFP in those regions on adjacent sections; R1-R2 denote ipsilateral tumor, R3-R5 denote tumor within corpus callosum, R6-R8 denote infiltrating tumor in contralateral hemisphere. **d.** Heatmap depicting the expression of markers associated with the single cell cluster of interest and their relative expression across the tumor regions displayed in **c**. Fold Change of markers from the scRNA-Seq data are mapped in pink and blue. **e.** Heatmap depicting the enrichment of markers associated with the single cell cluster of interest and their relative expression across various anatomical locations in human GBM, derived from the IVY-GAP database. These enrichment scores were generated with AUCell analysis (depicted in red and blue), or ssGSEA analysis (depicted in yellow and blue) and plotted as a heatmap

### Axon guidance genes drive glioma progression

The enrichment of axon guidance genes in the activity-driven, infiltrating glioma population (**Fig.3b**), led us to investigate their contributions to glioma infiltration. To examine their roles in this context, we performed a bar-coded, overexpression screen by generating a PiggyBac-based, barcoded library of 43 axon guidance-associated genes in our IUE-HGG model (**Fig.4a and Extended Table 5**). Following introduction of the axon guidance library, we harvested tumor bearing mice at P90 and dissected tumors based on ipsilateral- (primary) and contralateral- (secondary) locations, with the contralateral population likely enriched in infiltrating populations. Barcode sequencing was performed on samples from both sites, and we compared barcode enrichment between primary and secondary sites, seeking to identify those barcodes that are enriched in secondary sites, as those are candidate drivers of infiltration (**Fig.4a-b**). This analysis nominated a series of candidates enriched in secondary sites, including Unc5B^31,32^, Sema7A^33,34^, and Sema3C^35,36^, which have been previously implicated in tumor invasion (**Fig.4b**). Next, we evaluated the expression of genes enriched at secondary sites in the IVY-GAP database, finding that Semaphorin-4F (Sema4F), EphrinA6 (EphA6), and EphrinA7 (EphA7) are enriched in the leading edge of human GBM (**Extended Data Figure 4a-c**), while validating protein expression in primary GBM tumor samples (**Fig.4f**). Furthermore, the roles of EphA6, EphA7, and Sema4F in glioma infiltration are undefined, prompting us to further examine their contribution to tumorigenesis.

**Figure 4.**
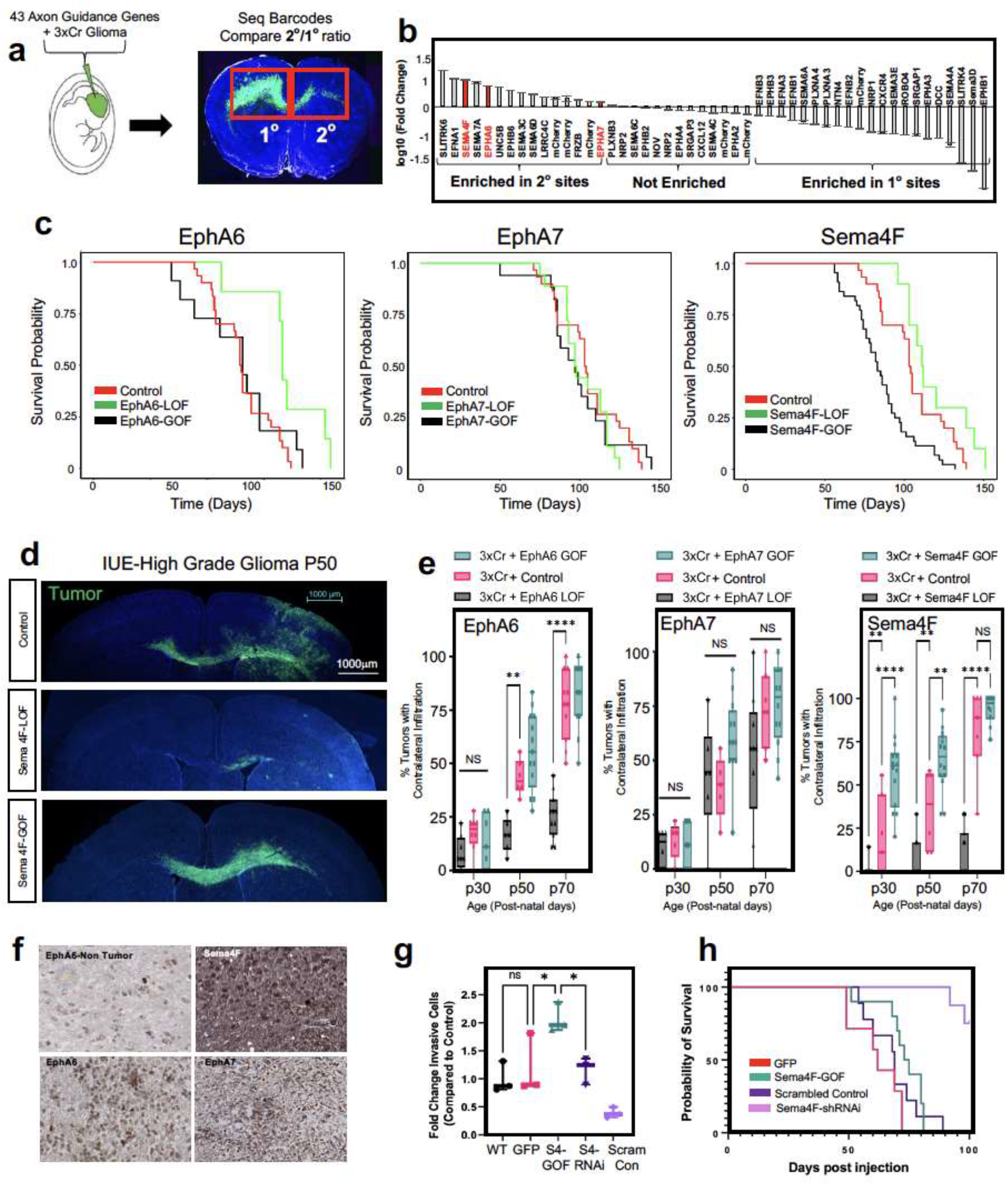
In vivo screen identifies Sema4F as a driver of glioma infiltration. **a.** Schematic of barcoded screen of 43 axon guidance genes and 7 internal mCherry controls. Tumors were harvested from ipsilateral primary and contralateral secondary tumor sites and barcode sequencing performed (n=4 tumors with paired sites). **b.** Next-generation sequencing for barcode amplification, fold-change was calculated with library input control and relative enrichment in primary or secondary site was determined. **c.** Kaplan-Meier survival curve of individual gain-of-function and loss-of-function validation studies for EphA6, EphA7 and Sema4F. EphA6-GOF (median_A6GOF_ = 105 days, Chisq = 0.7, df=1, p-value =0.4, n=11), EphA6-LOF (median_A6LOF_ = 133days, Chis =7.2, df=1, p-value=0.007, n = 7), EphA7-GOF (median_A7GOF_= 97 days, Chisq=0, df=1 p-value=0.9, n=15), EphA7-LOF (median_A7LOF_= 97.5 days, Chisq=1.3, df=1, p-value=0.3), Sema4F-GOF (median_S4FGOF_=83 days, Chisq = 14.8, df = 1 p-value = 0.0001, n = 44), Sema-LOF (median_S4FLOF_=112 days, Chisq = 3.9, df = 1 p-value = 0.05, n = 10), controls (n=18) **d.** Representative images from IUE-HGG tumors at P50 from Sema4F-GOF, Sema4F-LOF, or control groups demonstrating infiltration; green is tumor. **e.** Quantification of infiltration from these tumors across the P30-P70 timecourse. Infiltration was quantified based on the presence of tumor cells in contralateral cortex and analyzed via two-way analysis of variance (ANOVA). Error bars represent standard deviation, data derived from EphA6-GOF p30 n=7, EphA6-GOF p50 n=13, EphA6-GOF p70 n=11, EphA6 CTL p30 n=8, EphA6 CTL p50 n=6, EphA6 CTL p70 n=11, EphA6-LOF p30 n=8, EphA6-LOF p50 n=6, EphA6-LOF p70 n=11, EphA7-GOF p30 n=9, EphA7-GOF p50 n=12, EphA7-GOF p70 n=12, EphA7 CTL p30 n=5, EphA7 CTL p50 n=5, EphA7 CTL p70 n=7, EphA7-LOF p30 n=6, EphA7-LOF p50 n=6, EphA7-LOF p70 n=9, Sema4F-GOF p30 n=14, Sema4F-GOF p50 n=14, Sema4F-GOF p70 n=10, Sema4F CTL p30 n=7, Sema4F CTL p50 n=6 Sema4F CTL p70 n=7, Sema4F-LOF p30 n=17, Sema4F-LOF p50 n=16, Sema4F-LOF p70 n=19. **f.** Representative immunostainings of EphA6, EphA7, and Sema4F human tumor micro-array. **g.** Quantification of transwell migration of human glioma cell lines; infiltrating cells were counted after 48 hours incubation (n=3 wells per condition). **h.** Kaplan-Meier survival curve for human glioma cell lines transplanted into mouse brain. Samples were analyzed via log-rank (Mantel-Cox) test. WT median survival=66 days, n=11; GFP median=62 days n=7; Sema4f-GOF median=74 days, n=10, Chi = 0.4120, p-value = 0.5209; shSCR median=69 days, n=9; shSema4F median = undefined after 100 days, n=8, Chi = 14.08 p-value = 0.0002. \**P* < 0.05, \*\**P* < 0.01, \*\*\**P* < 0.001, **** *P* < 0.0001, two-way analysis of variance (ANOVA) (**e**), one-way analysis of variance (ANOVA) (**f**), Log-rank (Mantel-Cox) test (**c,g**)

To determine the roles of Sema4F, EphA6, and EphA7 in glioma tumorigenesis and infiltration, we performed gain-of-function (GOF) overexpression and CRISPR-Cas9 based loss-of-function (LOF) studies in our IUE-HGG model (**Extended Data Figure 4d**). Using overall survival as a proxy for tumor burden, we found that LOF studies with EphA6 and Sema4F extended mouse survival, while GOF studies with Sema4F decreased overall survival (**Fig.4c**). To determine how these manipulations impacted tumor infiltration we generated LOF and GOF tumors from each gene and harvested tumors at P30, P50, and P70, measuring contralateral infiltration. Consistent with our overall survival studies, we found that GOF manipulations with Sema4F accelerated infiltration, while LOF manipulations impaired infiltration (**Fig.4d-e**). Analysis of LOF of EphA6 revealed in impaired infiltration, while the remainder of the manipulations with EphA6 and EphA7 had relatively modest impacts on infiltration while retaining high-grade glioma histopathology (**Fig.4d-e and Extended Data Figure 5a-c**). Focusing on Sema4F, we generated human glioma cell lines that overexpress Sema4F (GOF) or have shRNA-knockdown of Sema4F (LOF) **(Extended Data Figure 5d**). We implanted these cell lines (and control cell lines) into the mouse brain and found that knockdown of Sema4F resulted in a significant extension of overall survival (**Fig.4h**). To evaluate glioma cell infiltration, we performed transwell assays, finding that Sema4F-GOF resulted in enhanced infiltration, while Sema4F-LOF suppressed infiltration (**Fig.4g**). Together, these studies indicate that our *in vivo* screening approach can identify new regulators of glioma infiltration, highlighting the role of Sema4F in glioma tumorigenesis.

### Sema4F is required for activity dependent infiltration

The central role of Sema4F in glioma infiltration raises the question of whether it is required for activity-dependent infiltration. To address this question, we used our activity-driven paradigm of glioma progression (**Fig.1a**), in combination with LOF of Sema4F. As before, we stimulated with CNO and employed saline controls starting at P20, harvesting tumor bearing brains at P30 and assessed contralateral infiltration, as well as Ki67 expression. Our analysis revealed that that loss of Sema4F in the context of CNO-based stimulation abolished activity-dependent infiltration when compared to controls containing Sema4F (**Fig.5a-b**). Additionally, Sema4F LOF abolished the effect of neural stimulation on tumor proliferation as measured by Ki67 staining, which demonstrated no significant differences between CNO and saline treated tumors (**Fig.5b and Extended Data Figure 6a-b**). Next, we examined whether the Sema4F contributes to glioma tumorigenesis via cell extrinsic mechanisms by overexpressing the Sema4F-ectodomain (S4E) in our IUE-HGG model, finding that it promotes infiltration and proliferation at P30 (**Extended Data Figure 6c-f**). These findings prompted us to examine whether S4E can rescue the deficits manifest in Sema4F-LOF tumors. Therefore, we overexpressed S4E in the context of Sema4F-LOF, finding that expression of S4E can rescue both infiltration and proliferation at P30 (**Extended Data Figure 6c-f**). Together, these results indicate that Sema4F is a key mediator of activity-dependent glioma and that it promotes tumor infiltration through its ectodomain, suggesting that it regulates this phenomenon through interactions with the brain microenvironment.

**Figure 5.**
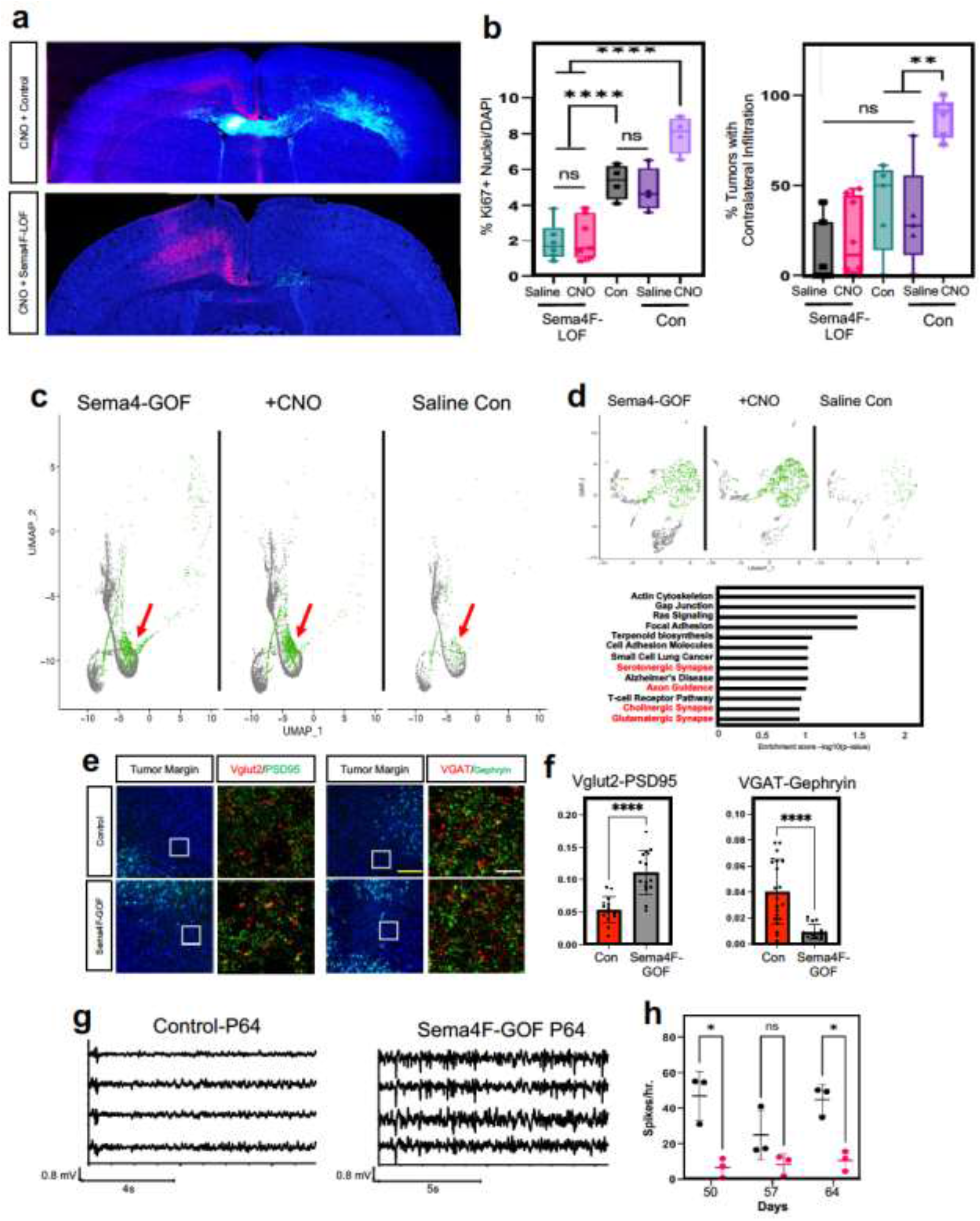
Sema4F promotes synaptic remodeling and brain hyperactivity. **a.** Representative images from IUE-HGG tumors at P30 demonstrating the extent of infiltration with combined Sema4F-LOF and neuronal activation (CNO) or neuronal activation control; green is tumor, red is AAV-DREADD virus. **b.** Quantification of tumor infiltration and Ki67 expression at P30; Infiltration was quantified based on the presence of tumor cells in contralateral cortex and analyzed via one-way analysis of variance (ANOVA). Error bars represent standard error, infiltration data derived from Sema4F-KO+AAV+Saline n=7, Sema4F-KO+AAV+CNO n=8, 3xCr CTL n=5, 3xCr+AAV+Saline n=5, 3xCr+AAV+CNO n=6. Proliferation data derived from Sema4F-KO+AAV+Saline n=6, Sema4F-KO+AAV+CNO n=7, 3xCr CTL n=4, 3xCr+AAV+Saline n=4, 3xCr+AAV+CNO n=4. **c**. Single Cell RNA-Seq DimPlots of P50 IUE-HGG from Sema4F-GOF, CNO, and saline controls. Cell types were mapped in SingleR with celldex as expression profile reference; shown is the GFP+ tumor cluster. Infiltrating tumor subpopulation is highlighted in green and denoted by red arrow. **d.** Sub-clustering analysis of tumor cells shown in GO-term analysis on the unique prospective cell population (in green) was performed using EnrichR and the KEGG 2021 Human dataset. **e.** Antibody staining of excitatory (Vglut2-PSD95) and inhibitory synapses (VGAT-Gephryin) P50 mouse brains at peritumoral margins from Sema4F and control tumors; box denotes zoomed in region in adjacent panel (10X and 200X magnification left to right; white scale bar is 12.5μm and yellow scale bar is 200μm). **f.** Quantification of synaptic staining derived from 3 separate tumors for each condition. Error bars represent standard error, data derived from 3 tumors derived from 3 mice with n≥15 fields analyzed and quantified, per condition. **g.** Sample EEG traces from mice bearing control or Sema4F-GOF tumors. **h.** Quantification of spikes/hr over a 24-hr period at one-week intervals from P50 to P64. Spikes were recorded in 3 mice per condition and analyzed via one-way analysis of variance (ANOVA). \**P* < 0.05, \*\**P* < 0.01, \*\*\**P* < 0.001, **** *P* < 0.0001, one-way analysis of variance (ANOVA) (**f,h**)

### Sema4F promotes synaptic remodeling and brain hyperactivity

The foregoing data also suggest that Sema4F itself promotes the generation of the activity-dependent, infiltrating population (**Fig.3**). To test this, we performed scRNA-Seq on Sema4F-GOF tumors at P50 and cross-compared these data with our activity-driven scRNA-Seq datasets (**Fig.5c and Extended Data Figure 7a**). This analysis revealed that the same cluster enriched in our activity-dependent dataset, was also enriched in our Sema4F-GOF dataset and contains the corresponding axon guidance gene signature (**Fig.5c-d; Extended Tables 2, 6**). These data suggest that Sema4F expression is capable of generating activity-dependent infiltrating glioma populations. Further analysis of our Sema4F-GOF scRNA-Seq dataset identified the upregulation of several synaptic signaling pathways in tumor cells **(Fig.5d**), including glutamatergic synapse genes. Bulk RNA-Seq of Sema4F-GOF human glioma cell lines revealed an analogous enrichment in synaptic signaling and axon guidance pathways, suggesting conserved function of Sema4F across these model systems (**Extended Figure 7c-d; Extended Tables 7-8**). Prior studies have shown that Sema-family members and their PlexinB receptors, which are expressed in neurons from our scRNA-Seq data (**Extended Data Figure 7b**), can engender synapse formation ^37^. These observations, coupled with the role of the Sema4F-ectodomain in tumorigenesis (**Extended Data Figure 6c-f**) prompted us to assess excitatory- and inhibitory-synapses in neurons outside the tumor margins in Sema4F-GOF tumors. These studies revealed a marked decrease in inhibitory synapses (VGAT-Gephryin), coupled with an increase in excitatory synapses (Vglut2-PSD95) in mouse Sema4F-GOF tumors (**Fig.5e-f**), which we also observed in mice bearing Sema4F-GOF tumors derived from human glioma cell lines (**Extended Data Figure 8**). Together these data indicate extensive synaptic remodeling towards hyperactive states. Next, we examined whether these alterations in the synaptic milieu influence brain network activity by performing serial electroencephalograms (EEG) on mice bearing Sema4F GOF tumors^15^. As shown in **Figure 5g-h,** mice bearing Sema4F-GOF tumors exhibit an early onset of brain network hyperactivity, featuring increased spiking, compared to mice bearing control tumors. These EEG data indicate that synaptic remodeling by Sema4F promotes brain network hyperactivity and in conjunction with the scRNA-Seq data suggests that Sema4F itself drives the generation of these activity dependent, infiltrating glioma populations.

## Discussion

Neuronal activity has emerged as a key component of the TME that engenders malignant growth in gliomas and across a host of cancers^15–21,38–40^, however the nature of tumor-neuron interactions remains incompletely understood. In this study, we use CPN activation to demonstrate that long-range projections from neuronal populations remote to primary glioma can drive progression and infiltration. Neuronal axons can extend across relatively long distances from their cell bodies. Therefore, our findings suggest that brain tumors receive inputs from a host of brain regions which implies a broader relationship between brain tumors and resident neurons than previously thought^28^. Glioma and neurons make direct synaptic connections^16^. Given our findings it is likely that circuit disruption is not limited to regions where the primary tumor resides but is more widespread throughout the brain. Furthermore, glioma tumors remodel local neuronal synapses towards hyperactivity^15–20^, raising the possibility that synapses from these long-range projections are also remodeled by the tumor resulting in deleterious effects on brain circuits in remote regions. Interestingly, recent studies from mouse glioma models revealed spreading depolarization and hyperactivity across cortical hemispheres^41^, suggesting dysregulation of circuits remote to the primary tumor.

Despite its central role in glioma recurrence, the cellular and molecular mechanisms regulating tumor infiltration remain elusive. We identified an activity-dependent infiltrating glioma population, which indicates that glioma tumors utilize neuronal signals to drive progression and widespread infiltration. The precocious emergence of this population is facilitated by neuronal activity, however our identification of this population in Sema4F-GOF tumors and at later stages of progression suggest that its emergence is a core feature of infiltration. Glioma tends to use white matter tracts as routes of infiltration. These myelinated axonal structures are populated by nodes of Ranvier, which are sources of dynamic ion flux during activity and could serve as an infiltrative cue^9,11,42–44^. Mechanistically, we found that the infiltrating population is enriched for axon guidance genes and our *in vivo* screen identified Sema4F as a key driver of glioma progression and activity-dependent infiltration. Despite playing key roles in responding to environmental cues during development, roles for axon guidance genes in glioma remain poorly defined and our studies highlight their central role in activity-dependent glioma progression^45–47^. An intriguing line of future investigation is to decipher how Ephrin- and Sema-family members cooperate to regulate glioma infiltration and progression. Further analysis revealed that Sema4F promotes synaptic remodeling in neurons adjacent to glioma, which is consistent with prior models suggesting that tumors in the CNS generate a positive feedback loop of receiving and promoting synaptic signaling to tumor populations^15–17,22,41^. When put together, a model emerges where neurons provoke expression of genes from glioma tumors that subsequently drive their own synaptic activity.

## Acknowledgements

This work was supported by US National Institutes of Health grants NS124093, NS071153, and CA223388 to BD. This work was also supported by the National Cancer Institute-Cancer Target Discovery and Development, U01-CA217842 to BD. In addition, F31-CA243382 to E.H.H, 1F31CA265156 to RNC, T32-5T32HL092332-19 to BL, and and NIH Director’s Pioneer Award DP1NS111132 to M.M. We are thankful for support from the David and Eula Wintermann Foundation. scRNA-Seq studies were performed at the Single Cell Genomics Core at BCM partially supported by NIH shared instrument grants (S10OD023469, S10OD025240) and P30EY002520. Human tumor tissue samples were obtained from the Dan L. Duncan Cancer Center Pathology and Histology Core (HTAP) core at Baylor College of Medicine (IRB#: H-35355), supported by P30 Cancer Center Support Grant (NCI-CA125123). We would like to acknowledge the Optogenetics and Viral Vectors Core at the Jan and Dan Duncan Neurological Research Institute. Research reported in this publication was supported by the Eunice Kennedy Shriver National Institute of Child Health & Human Development of the National Institutes of Health under Award Number P50HD103555 for use of the Microscopy Core facilities and the Animal Phenotyping & Preclinical Endpoints Core facilities.

## Authors Contributions

EHH and BD conceived the project and designed the experiments; EHH, YTC, YK, ELF, YY, KRT, MMc, PH, HCC, EM, ZFL, SM, MW, and DC performed the experiments; JW executed the electrophysiology studies; RNC, MM, AJ, JLN, GR provided essential reagents; EHH, BL, ASH, and JB designed and executed the bioinformatics analyses. KRT and MM designed and performed in vitro glioma migration experiments. EHH and BD wrote the manuscript.

## References

1. Louis, D. N. et al. The 2021 WHO Classification of Tumors of the Central Nervous System: a summary. Neuro. Oncol. 23, 1231–1251 (2021).

2. Wen, P. Y. & Kesari, S. Malignant Gliomas in Adults. N. Engl. J. Med. 359, 492–507 (2008).

3. Omuro, A. & LM, D. Glioblastoma and other malignant gliomas: A clinical review. JAMA 310, 1842–1850 (2013).

4. Weller, M. et al. Glioma. Nat. Rev. Dis. Prim. 1, 15017 (2015).

5. Konishi, Y., Muragaki, Y., Iseki, H., Mitsuhashi, N. & Okada, Y. Patterns of Intracranial Glioblastoma Recurrence After Aggressive Surgical Resection and Adjuvant Management: Retrospective Analysis of 43 Cases. Neurol. Med. Chir. (Tokyo). 52, 577–586 (2012).

6. Milano, M. T. et al. Patterns and Timing of Recurrence After Temozolomide-Based Chemoradiation for Glioblastoma. Int. J. Radiat. Oncol. 78, 1147–1155 (2010).

7. McDonald, M. W., Shu, H.-K. G., Curran, W. J. & Crocker, I. R. Pattern of Failure After Limited Margin Radiotherapy and Temozolomide for Glioblastoma. Int. J. Radiat. Oncol. 79, 130–136 (2011).

8. Maher, E. A. & Bachoo, R. M. Glioblastoma. Rosenberg’s Molecular and Genetic Basis of Neurological and Psychiatric Disease: Fifth Edition (2014). doi:10.1016/B978-0-12-410529-4.00078-4.

9. Vollmann-Zwerenz, A., Leidgens, V., Feliciello, G., Klein, C. A. & Hau, P. Tumor Cell Invasion in Glioblastoma. Int. J. Mol. Sci. 21, 1932 (2020).

10. Vitorino, P. & Meyer, T. Modular control of endothelial sheet migration. Genes Dev. 22, 3268–3281 (2008).

11. Cuddapah, V. A., Robel, S., Watkins, S. & Sontheimer, H. A neurocentric perspective on glioma invasion. Nat Rev Neurosci 15, 455–465 (2014).

12. Numan, T. et al. Non-invasively measured brain activity and radiological progression in diffuse glioma. Sci. Rep. 11, 18990 (2021).

13. Robert, S. M. et al. SLC7A11 expression is associated with seizures and predicts poor survival in patients with malignant glioma. Sci. Transl. Med. 7, 289ra86 (2015).

14. Buckingham, S. C. et al. Glutamate release by primary brain tumors induces epileptic activity. Nat. Med. 17, 1269–1274 (2011).

15. Yu, K. et al. PIK3CA variants selectively initiate brain hyperactivity during gliomagenesis. Nature 578, (2020).

16. Venkatesh, H. S. et al. Electrical and synaptic integration of glioma into neural circuits. Nature 573, 539–545 (2019).

17. Venkatesh, H. S. et al. Neuronal Activity Promotes Glioma Growth through Neuroligin-3 Secretion. Cell 161, 803–816 (2015).

18. Pan, Y. et al. NF1 mutation drives neuronal activity-dependent initiation of optic glioma. Nature 594, 277–282 (2021).

19. Chen, P. et al. Olfactory sensory experience regulates gliomagenesis via neuronal IGF1. Nature 606, 550–556 (2022).

20. Venkataramani, V. et al. Glutamatergic synaptic input to glioma cells drives brain tumour progression. Nature 573, 532–538 (2019).

21. Venkataramani, V. et al. Glioblastoma hijacks neuronal mechanisms for brain invasion. Cell 185, 2899–2917.e31 (2022).

22. Venkatesh, H. S. et al. Targeting neuronal activity-regulated neuroligin-3 dependency in high-grade glioma. Nature 549, 533–537 (2017).

23. Doucette, T. et al. Bcl-2 promotes malignant progression in a PDGF-B-dependent murine model of oligodendroglioma. Int. J. cancer 129, 2093–2103 (2011).

24. Holland, E. C. & Varmus, H. E. Basic fibroblast growth factor induces cell migration and proliferation after glia-specific gene transfer in mice. Proc. Natl. Acad. Sci. U. S. A. 95, 1218–1223 (1998).

25. Harpold, H., Ellsworth, CA, & Swanson K.R. The evolution of mathematical modeling of glioma proliferation and invasion. J Neuropathol Exp Neurol. 66, 1–9 (2007)

26. Nagaraja, S. et al. Transcriptional Dependencies in Diffuse Intrinsic Pontine Glioma. Cancer Cell 31, 635–652.e6 (2017).

27. Vinci, M., Box, C., Zimmermann, M. & Eccles, S. A. Tumor spheroid-based migration assays for evaluation of therapeutic agents. Methods Mol. Biol. 986, 253–266 (2013).

28. Fame, R. M., MacDonald, J. L. & Macklis, J. D. Development, specification, and diversity of callosal projection neurons. Trends Neurosci. 34, 41–50 (2011).

29. Harris, J. A. et al. Anatomical characterization of Cre driver mice for neural circuit mapping and manipulation. Front. Neural Circuits 8, 76 (2014).

30. Puchalski, R. B. et al. An anatomic transcriptional atlas of human glioblastoma. Science (80-.). 360, 660–663 (2018).

31. Durko, M. et al. Rat C6 glioma cell motility and glioma growth are regulated by netrin and netrin receptors unc5B and DCC. J. Cancer Ther. Res. 2, (2013).

32. Wu, S. et al. High expression of UNC5B enhances tumor proliferation, increases metastasis, and worsens prognosis in breast cancer. Aging (Albany. NY). 12, 17079–17098 (2020).

33. Formolo, C. A. et al. Secretome Signature of Invasive Glioblastoma Multiforme. J. Proteome Res. 10, 3149–3159 (2011).

34. Manini, I. et al. Semaphorin-7A on Exosomes: A Promigratory Signal in the Glioma Microenvironment. Cancers vol. 11 at https://doi.org/10.3390/cancers11060758 (2019).

35. Vaitkienė, P. et al. High level of Sema3C is associated with glioma malignancy. Diagn. Pathol. 10, 58 (2015).

36. Yin, L. et al. MAOA promotes prostate cancer cell perineural invasion through SEMA3C/PlexinA2/NRP1–cMET signaling. Oncogene 40, 1362–1374 (2021).

37. McDermott, J. E., Goldblatt, D. & Paradis, S. Class 4 Semaphorins and Plexin-B receptors regulate GABAergic and glutamatergic synapse development in the mammalian hippocampus. Mol. Cell. Neurosci. 92, 50–66 (2018).

38. Ayala, G. E. et al. Cancer-Related Axonogenesis and Neurogenesis in Prostate Cancer. Clin. Cancer Res. 14, 7593–7603 (2008).

39. Li, J., Kang, R. & Tang, D. Cellular and molecular mechanisms of perineural invasion of pancreatic ductal adenocarcinoma. Cancer Commun. 41, 642–660 (2021).

40. Claire, M. et al. Autonomic Nerve Development Contributes to Prostate Cancer Progression. Science (80-.). 341, 1236361 (2013).

41. Hatcher, A. et al. Pathogenesis of peritumoral hyperexcitability in an immunocompetent CRISPR-based glioblastoma model. J. Clin. Invest. 130, 2286–2300 (2020).

42. Rasband, M. N. & Peles, E. Mechanisms of node of Ranvier assembly. Nat. Rev. Neurosci. 22, 7–20 (2021).

43. Brandalise, F. et al. Deeper and Deeper on the Role of BK and Kir4.1 Channels in Glioblastoma Invasiveness: A Novel Summative Mechanism?. Frontiers in Neuroscience vol. 14 at (2020).

44. Seker-Polat, F., Pinarbasi Degirmenci, N., Solaroglu, I. & Bagci-Onder, T. Tumor Cell Infiltration into the Brain in Glioblastoma: From Mechanisms to Clinical Perspectives. Cancers (Basel). 14, 443 (2022).

45. Li, X., Law, J. W. S. & Lee, A. Y. W. Semaphorin 5A and plexin-B3 regulate human glioma cell motility and morphology through Rac1 and the actin cytoskeleton. Oncogene 31, 595–610 (2012).

46. Le, A. P. et al. Plexin-B2 promotes invasive growth of malignant glioma. Oncotarget 6, 7293–7304 (2015).

47. Wang, L.-F. et al. Increased expression of EphA7 correlates with adverse outcome in primary and recurrent glioblastoma multiforme patients. BMC Cancer 8, 79 (2008).

48. Mascheroni P, et al. On the impact of chemo-mechanically induced phenotypic transitions in glioma. Cancers. 11, 716 (2019)

