## Supplementary material for "Remote neuronal activity drives glioma infiltration via Sema4f": Methods and Extended Data Figures 1-8

In utero electroporation model of HGG, RCAS model of LGG

All mouse gliomas were generated in the CD-1 IGS mouse background as previously described (Yu, et al. Nature 2020). In utero electroporation was performed on embryonic day 15. Previously generated CRISPR constructs were used to knockout NF1, PTEN, and TRP53 (each at 1.5µg per µl) (Yu, et al Nature 2020). For our low grade glioma studies we used the RCAS/Ntv-a system, a mouse model of platelet-derived growth factor subunit B (PDGFB)–driven gliomas. To generate RCAS/PDGFB virus we used immortalized DF-1 chicken fibroblasts grown in Dulbecco modified Eagle medium containing 10% fetal bovine serum in a humidified atmosphere (95% air and 5% carbon dioxide) at 37 C. Live virus was produced by transfecting the RCAS/PDGFB vector into DF-1 cells using FuGENE-6 (Roche, Indianapolis, IN) and allowing the cells to replicate in culture. DF-1, viral producing cells were then injected into the cortex of P1 RCAS/Ntv-a pups. All procedures were approved by the Institutional Animal Care and Use Committee at Baylor College of Medicine and conform to the US Public Health Service Policy on Human Care and Use of Laboratory Animals.

Plasmid and adeno-associated virus (AAV) generation

For the initial contralateral neural stimulation experiments, pAAV-hSyn-hM3D-2a-mCherry (Addgene plasmid #50474) was used to generate AAV 2/9 at a concentration of 1.73E13 gc/mL genome copies per mL (gc/mL). Additional experiments in Rasgrf2-dCre mice utilized pAAV-DIO-EF1a-hM3D-mCherry (Addgene plasmid #50460) to generate AAV2/9 at a concentration of 1.02E13 gc/mL. pAAV-GqDREADD-P2A-dTomato-Fishell-

4 plasmid (Addgene #83897) was used to generate AAV 2/9 to target GABA-ergic interneurons. Virus was produced at 3.14e11 gc/mL. AAV-DREADDs were generated at the Neuroconnectivity Core at Baylor College of Medicine.

AAV-DREADD injection and CNO treatment

AAV was diluted with loading dye (10x Fast Green 2mg/mL) loaded into a micro dispenser (Drummond Scientific, Cat no.13-681-460) and injected at rate of 7nL/sec to a total volume of 1 µL into the left ventricle of 5-7 day old mice. After weaning, mice were weighed daily and injected intra-peritoneally with 1mg/mL Clozapine N-oxide (CNO) (Tocris Cat #4936) at 0.5 mg/kg per mouse or an equivalent volume of Saline for controls. Mice received injections of CNO or Saline respectively twice per day for the duration of their respective studies reaching 30, 50, or 70 days of age.

Histological Sample Preparation

Mice were anesthetized under isofluorane inhalation and perfused transcardially with ice-cold 1XPBS pH 7.4 followed by 10% Formalin for paraffin embedded samples, or 4% PFA for frozen sections. After fixing overnight on a shaker at 4C, samples were moved to 70% ethanol or 40% Sucrose respectively. Paraformaldehyde/Sucrose fixed samples were then embedded in OCT (Sakura Finetek, 4583) after an additional overnight incubation on a shaker at 4C. Formalin/Ethanol fixed samples were submitted to the Pathology Core and Lab at Baylor College of Medicine for paraffin embedding.

For haematoxylin and eosin (H&E) staining, 10-μm paraffin-embedded sections were process as follows: 3 × 3 min in xylene, 3 × 3 min in 100% ethanol, 3 × 3 min in 95% ethanol, 3 min in 80% ethanol, 5 min in 70% ethanol, 5 min in dd$H_2O$, 2.5 min in Harris haematoxylin (Poly Scientific R&D, S212A), running tap water wash, 30 s in 95% ethanol, 2.5 min in eosin (Poly Scientific R&D, S176), 2 × 2 min in 95% ethanol, 2 × 2 min in 100% ethanol, and 2 × 2 min in xylene. Staining was preserved with Permount Mounting Media (Electron Microscope Sciences, 17986-01) under a coverslip. Histological diagnoses of mouse-IUE-generated tumors as well as RCAS-Ntva tumors were validated across $n \geq 4$ tumors per variant. FFPE sections were subject to antigen retrieval (when needed), blocking and staining with the following antibodies: EphrinA6 (LS Bio LS-B4905, 1:250), EphrinA7(LSBio LS-B4098, 1:500), Sema4F (Atlas Antibodies HPA064095, 1:500), Ki67 (abcam #16667, 1:250) overnight at 4 °C. Slides were washed the following morning in 1xPBS 5 min X3 and stained with secondary antibody (ImmPRESS (Peroxidase) Polymer Anti-Rabbit IgG Reagent) at room temperature for 1 hour then stained with DAB (3,3'-diaminobenzidine) peroxidase substrate (Vector Labs, SK4100). Nuclei were counterstained with Hematoxyline for 1 minute followed by development in running water for 5 minutes. Slides were dehydrated with 1x 50% ethanol 5min, 1x 75% ethanol 5min, 3x 95% ethanol 5 min, 3x xylene 5 min followed by mounting with Permount (Fisher chemical SP15). A 20 sided die was used to randomly sample RCAS tumor slides from each respective brain.

Frozen brains were sectioned at 20 μm thickness. Sections were subject to antigen retrieval (when needed), blocking and staining with Ki67 (abcam #16667, 1:250) overnight at 4 °C. We used rabbit-specific secondary antibodies tagged with Alexa Fluor 647

(1:1,000, ThermoFisher) for immunofluorescence. After Hoechst nuclear counter staining (ThermoFisher, H3570, 1:50,000), coverslips were mounted with VECTASHIELD antifade mounting medium (Vector Laboratories, H-1000).

Image Analysis of Ki67+ Nuclei

Ki67 staining images were taken using a Zeiss Axio Imager M2 microscope. Ki67 positive nuclei and total nuclei number were recorded for each field of view in ImageJ/Fiji and the percentage calculated. For RCAS slides utilizing DAB staining, images were taken at 20X magnification and analyzed with ImageJ/Fiji. For IUE-HGG slides utilizing Alexa-Fluor 647, 20x images were taken and analyzed with ImageJ/Fiji.

Infiltration Analysis

Tumor brains were harvested and coronal sections containing GFP positive tumor were collected. From coronal sections containing GFP positive tumor, cortex contralateral to the tumor origin was observed and the presence of GFP tumor cells was logged. Sample brains were sampled randomly with N ≥ 3 slides randomly sampled from regions with GFP. Each slide contained at least N ≥ 3 coronal sections per slide. Infiltration in the contralateral cortex was converted to a percentage for each individual brain. Resulting percentages were analyzed and compared via one-way or two-way ANOVA in Prism (v9.3.0).

Corpus Callosum Cutting Surgery

After receiving viral injections at 5 days of age as outlined above, pups were anesthetized at 7-10 days of age with ice. Using a pair of fine foreceps (FST 11412-11), the skull was penetrated and corpus callosum severed. Penetrating ~2mm in depth and severing the corpus callosum between the 1-2 mm rostral to the bregma. Mice were sutured and monitored during recovery.

Rasgrf2-dCre Induction

Mice were injected at weaning with trimethoprim (Sigma-Aldrich – T7883) suspended in DMSO at 100mg/mL. Mice were injected intraperitoneally before weaning to a final concentration of 100ng/gram body weight[19]. As a control, Rasgrf2-dCre and Rosa26-LSL-TdTomato mice were crossed. 48 hours post-injection of trimethoprim, mice were sacrificed and analyzed for TdTomato expression in coronal slices (**Extended Data Figure 4)**. Seeing adequate activation, IUE tumors were generated and AAV injected as outlined above. Mice were injected 2 days before weaning with Trimethorpim and received CNO or Saline treatment for 10 days at weaning.

Single Cell Sequencing

Tumor bearing mice were anesthetized under isofluorane inhalation. Brains were isolated and dissected based on the presence of GFP (Leica M80 stereomicroscope, Leica EL6000 Fluorescence Light source). One male and one female mouse was isolated for each group to control for sex-dependent variability. GFP positive tumor tissue was isolated and placed on ice. Tumors were dissociated by incubating with papain (Worthington, LK003178) with DNAseI (Worthington, LK003172) resuspended in EBSS

(ThermoFisher, 24010043) on a Thermoshaker (Eppendorf, Z605271) for 15 minutes at 1400 rpm. Papain was neutralized using 10% FBS DMEM F12 w/PS and samples were spun down at 300g for 5 minutes. Samples were washed with PBS and RBC removal was performed as described (Miltenyi 130-094-183). Debris Removal was performed as described (Miltenyi 130-109-398). Following this we performed live cell cleanup using Dead Cell Removal

Kit (Miltenyi 130-090-101) and sequenced samples using 10X Genomics through the Baylor Single Cell Sequencing Core. Samples were analyzed using Seurat, AUCell, ssGSEA, and EnrichR.

Single cell mouse signature in patient leading edge tumor

Seurat R package is used for standard procedures for downstream analysis such as filtering, mitochondrial gene removal, variable gene selection, dimensionality reduction, and clustering. Cells are flagged for downstream analysis using the following criteria: mitochondrial RNA read content > 5%, number of genes detected in each cell (*nFeature_RNA*) > 5000 and < 200. The read count matrix is scaled/normalized and variance stabilization is used to eliminate technical noise from the data. Genes with the highest variation among cells are selected and reported. Multiple datasets are integrated using Harmony batch effect correction algorithm (**Korsunsky et al. 2019**). Dimensionality reduction is performed using UMAP, and PCA. The clustering is performed using dimensionality reduced representation. Seurat R package FindMarkers function was used using default settings to identify all differentially expressed markers between Leading Edge (LE) vs Core (Core) samples.

**Mathematical dissection of infiltration and proliferation**

To understand the dynamics of the glioma tumor cells we estimated two quantities namely infiltration width (IW) and tumor mass (TM) within each condition (CNO, Saline and Control) at the p30 time point. Images were taken at tumor core, contralateral infiltration, and midline points. Piecewise-cubic splines are used for interpolating and smoothing the data points. Using these smoothed values, within each condition the 80% (0.8 $p_{max}$) and 2% (0.02 $p_{max}$) glioma cell density of the maximum smoothed cellular density ($p_{max}$ ) was estimated. IW was defined by the difference between distance to the tumor origin at which glioma cell density is in 0.8 $p_{max}$ and 0.02 $p_{max}$ (25,48). TM was estimated using the areas under the smoothed curve.

Spatial Transcriptomics

GeoMx Digital Spatial Profiling (DSP) was performed through Nanostring. FFPE samples were sliced at 5µm thickness and stained for Vimentin (Santa Cruz - sc-373717), GFAP (Thermo - 53-9892-82), and Iba1 (Millipore - MABN92-AF647). Tumor cells were identified by co-expression of Vimentin and GFAP and no Iba1 expression. Adjacent sections were also stained for GFP (GeneTex - GTX113617) to confirm the presence of tumor. Next-generation sequencing was performed at Nanostring and data generated using established methods[26]. Data were subsequently analyzed and heatmaps generated in RStudio.

Barcode Screen

Barcoded axon guidance genes of interest were generated as previously described[27]. The list of these genes can be found in **Supplementary Table 3.** Pooled injections were performed as previously described[27]. Mice were euthanized at 90 days after demonstrating symptoms and tumor was isolated with fluorescence assisted microdissection of tumor as described above. Tumor from primary tumor sites and contralateral tumor sites was isolated and sequenced separately. Samples were prepared in biological and technical replicates (n = 4 biological replicates, n = 2 technical replicates) and compared to library input control (n = 2 technical replicates) to calculate fold change. Barcode libraries were prepared as previously reported[27,28]. Barcode read representation was calculated by quantifying the reads of each barcode divided by total barcode reads per sample. Barcode enrichment was calculated as fold change value relative to the library input control. Standard error of this fold change was calculated and plotted on the barcode enrichment graphs. A control average and standard error was calculated between mCherry-barcode constructs included in the library. These were used as a cutoff for significance and are plotted as red lines on the barcode enrichment graphs.

In vivo functionalization – Individual ORFs

In vivo functionalization of EPHA6, EPHA7 and SEMA4F was performed by co-electroporating a pBCAG construct containing orfs for each respective gene (EPHA6 – NM_173655, EPHA7 - NM_004440, SEMA4F - NM_004263) at 1.0 µg per µL final concentration in the established IUE cocktail. Loss of function studies were performed by co-electroporating a CRISPR construct containing guides targeting each respective gene.

The gRNA sequences were as follows: EPHA6 – 5’ GACAGGGTATGAAGAATCGA 3’, EPHA7 – 5’ GCACCTGGTATGTTCGTATCG 3’, SEMA4F - 5’ GACTACCTGTCATGGACAACG 3’. On and off target analysis was performed using Surveyor® Mutation Detection Kit (Integrated DNA Technologies; 706025) following manufacturer’s protocol. Western blots to confirm manipulations of tumors were performed using the following primary antibodies: EPHA6, 1:100 – Abcam: ab80207; EPHA7, Abcam: ab136095 ; SEMA4F 1:500 – LSBio: LS-C497142; Actin 1:500 – GeneTex: GTX629630. We used species specific secondary HRP conjugated antibodies from Jackson ImmunoResearch Labs to visualize western tagged blots.

Confocal Imaging

Tiling images were generated on a Zeiss LSM 880 laser scanning confocal microscope at 10X magnification. Coronal sections were scanned and tiled using Zeiss Zen Blue Edition software. Synaptic staining was imaged using a Zeiss LSM 880 laser scanning confocal microscope with 63X oil immersion. Colocalization of signal was analyzed using Synapse Counter (SynPuCo) in ImageJ/Fiji (https://github.com/SynPuCo/SynapseCounter)

EEG methods

Similarly to previous studies (Yu, et al. 2020, Hatcher, et al. 2020), IUE mice at P40 were anesthetized with isoflurane vaporization pump and surgically implanted with bilateral silver wire electrodes (0.005-in. diameter) attached to a microminiature connector. Electrodes were placed bilaterally through burr holes subdurally in the frontal and parietal lobes with two grounds placed in the cerebellum. Mice were recorded for 48 hours continuously every 7 days starting at post-natal day 50. EEG signals were filtered and analyzed in LabChart Pro 8 (ADI Instruments). EEG signals were filtered with a 1-50Hz band-pass filter and interictal spikes were quantified using the Peak Analysis tool in LabChart Pro and standardized per hour of recording.

**PDX methods:**

Patient-derived primary GBM cell lines were maintained in neurosphere media (DMEM/F12 (ThermoFisher, 11320082), supplemented with B27 (1X, ThermoFisher, 17504001), bFGF (20ng/mL, Peprotech, 100–18B), and EGF (20ng/mL, Peprotech, 100–47)). Cells were lentivirally infected with Sema4F overexpression, GFP control, shRNA constructs or shScramble control. After 48 hours infection, cells were selected for by puromycin (1μg/mL, ThermoFisher, A1113803). 50,000 live cells were stereotaxically transplanted into 6 week old female nude mouse (Taconic, CrTac:NCr-*Foxn1*$^{nu}$) brains (from bregma, cells were injected 1.0mm caudal, 2.0mm lateral, 2.5mm ventral).

<u>Human Cell line generation</u>

Human PDX lines were infected with lentivirus generated as previously described. shRNAi constructs were acquired from OriGene (TL309556). Over expression constructs were generated via gateway cloning into pLenti-GTWY-ires-Puro constructs . Cells were lentivirally infected and were selected with puromycin (1ug/mL ThermoFisher, A1113803) after 48 hours. Expression was validated via qPCR (**Extended Data Figure 5d**). RNA was isolated via RNeasy Plus Mini Kit (Qiagen, 74134). For RT-qPCR, 500 ng of RNA was converted to complementary DNA (cDNA) using iScript Reverse Transcriptase Supermix. (BioRad, 1708841). RT-qPCR was performed using PerfeCTa SYBR Green Fast Mix (Quantabio, 95072-012) on a Roche Light Cycler 480 instrument. Reactions were set up using 2 ng cDNA, 250 nM primers, and 1× SYBR mix. qPCR was carried out at 95 °C for 30 s, 40 cycles of 95 °C for 5 s and 60 °C for 30 s, with subsequent melting curve analysis. The expression of transcripts of target genes was normalized to *Gapdh*. Primers for qPCR were: Sema4F-CDS-F (CAGACCTGGAGAGTGCATCA), Sema4F-CDS-R (GCCACGACTCTGAGATAGGC), Gapdh-CTL-F (ACAACTTTGGTATCGTGGAAGG), Gapdh-CTL-R (GCCATCACGCCACAGTTTC).

<u>In vitro spheroid proliferation and infiltration</u>

For the cell invasion assay, differences in the cell invasive potential were assessed using the CytoSelect Cell Invasion Assay (Cell Biolabs, USA) according to the manufacturer's protocol with some modifications. 1.0x $10^5$ cells of each line were seeded in DMEM-F12 + 1% P/S into the basement membrane inserts. Inserts were placed in wells with DMEM + 10% FBS + 1% P/S. After 48 hours, invading cells were centrifuged, resuspended in 10uL PBS and counted by automated cell-counter (Bio-Rad).

<u>Three-dimensional (3D) migration and invasion assays</u>

3D migration assays were adapted from methods previously described (Vinci *et al*, 2013), with additional modifications. Briefly, cells were seeded at $10^3$/200ul/well in ULA 96-well round-bottomed plates and incubated for four days at 37°C, 5% $CO_2$, 95% humidity to facilitate spheroid formation. Flat-bottomed 96-well plates (Corning biosciences) were coated with 125μg/ml matrigel in stem-cell media. Once coating was completed, 200μl/well of culture medium containing either growth factor media, ACSF or conditioned media collected from acute cortical slices was added to each well. After removal of 100μl medium from the ULA 96-well round-bottomed plates containing neurospheres of 250-300μm in diameter, the remaining medium including the neurosphere was transferred onto the pre-coated plates (4 replicates). Images were collected using standard exposure and gamma settings starting from time zero, and at 24-hours. The degree of cell spread on the matrix was measured and data plotted as the average area of migration using Image J.

Conditioned media

Mice expressing Thy1::Chr2 (channelrhodopsin-expressing) were used at 4-7 weeks of age. Brief exposure to isoflurane rendered the mice unconscious prior to decapitation. ExtrBrains were extracted and placed in an oxygenated sucrose cutting solution and sliced at 350um as described previously (Venkatesh 2015). The slices were placed in ACSF (containing in mM 125 NaCl, 2.5 KCl, 25 glucose, 25 $NaHCO_3$, 1.25 $NaH_2PO_4$, 1 $MgCl_2$ and $CaCl_2$) and allowed to recover for 30 minutes at 37ºC and 30mins at room temperature. After recovery the slices were moved to fresh ACSF and stimulated using a blue-light LED using a microscope objective. The stimulation paradigm was 20-Hz pulses of blue light for 30 seconds on, 90 seconds off over a period of 30 minutes. The surrounding conditioned medium was collected and used immediately or frozen at -80ºC for future use.

Human cell line RNA-Seq

Cell lines were generated as described above. Cell pellets were collected and RNA was isolated using the RNeasy Plus Mini Kit (Qiagen, 74134) according to the manufacturer's protocol. Samples were prepared in biological replicates, N = 3 for each experimental group. RNA integrity (RIN ≥ 8.0) was confirmed using the High Sensitivity RNA Analysis Kit (AATI, DNF-472–0500) on a 12-Capillary Fragment Analyzer. Illumina sequencing libraries with 6-bp single indices were constructed from 1 μg total RNA using the TruSeq Stranded mRNA LT kit (Illumina, RS-122–2101). The resulting library was validated using the Standard Sensitivity NGS Fragment Analysis Kit (AATI, DNF-473–0500) on a 12-Capillary Fragment Analyzer. Equal concentrations (2nM) of libraries were pooled and subjected to sequencing of approximately 10-20 million reads per sample using the Mid Output v2 kit (Illumina, FC-404–2001) on a Illumina NextSeq550 following the manufacturer's instructions.

Sequencing files in fastq format were downloaded, and files from each flow cell lane were merged, followed by quality control analysis using fastQC (v0.10.1) and MultiQC (v0.9). Reads were aligned to the human genome using hg19 assembly by STAR (v2.5.0a). Mapped reads were used to build count matrices and gene models using Rsamtools (v2.0.0) and GenomicFeatures (v1.32.2) for expression quantification. UCSC transcripts were downloaded from Illumina iGenomes as GTF files. Reads per million were determined using GenomicAlignments (v1.16.0). DESeq2 was used for differential gene expression analysis and read count normalization. Plots for data visualization were generated using ggplot2 (v3.3.2).

Quantification and Statistical Analysis

Sample sizes and statistical tests can be found in figure legends. Analysis was carried out using Fiji, QuPath-0.3.2, Graphpad Prism 9, Excel 2016, and R-studio. For survival data analysis we used log-rank test and measured significance with Chi-Squared test statistic. For infiltration analysis, we performed one-way analysis of variance (ANOVA) with Tukey's test, or two-way analysis of variance (ANOVA) with Sidak's test of treatment groups based on infiltration into the cortex. For multiple comparisons, we used the one-way ANOVA with Tukey's test and two-way ANOVA with Sidak's test. In general, we assumed data were normally distributed but this was not formally tested. Data are presented as mean ± SEM (standard error of the mean). Levels of statistical significance are indicated as follows: * ($p < 0.05$), ** ($p < 0.01$), *** ($p < 0.001$), **** ($p < 0.0001$).

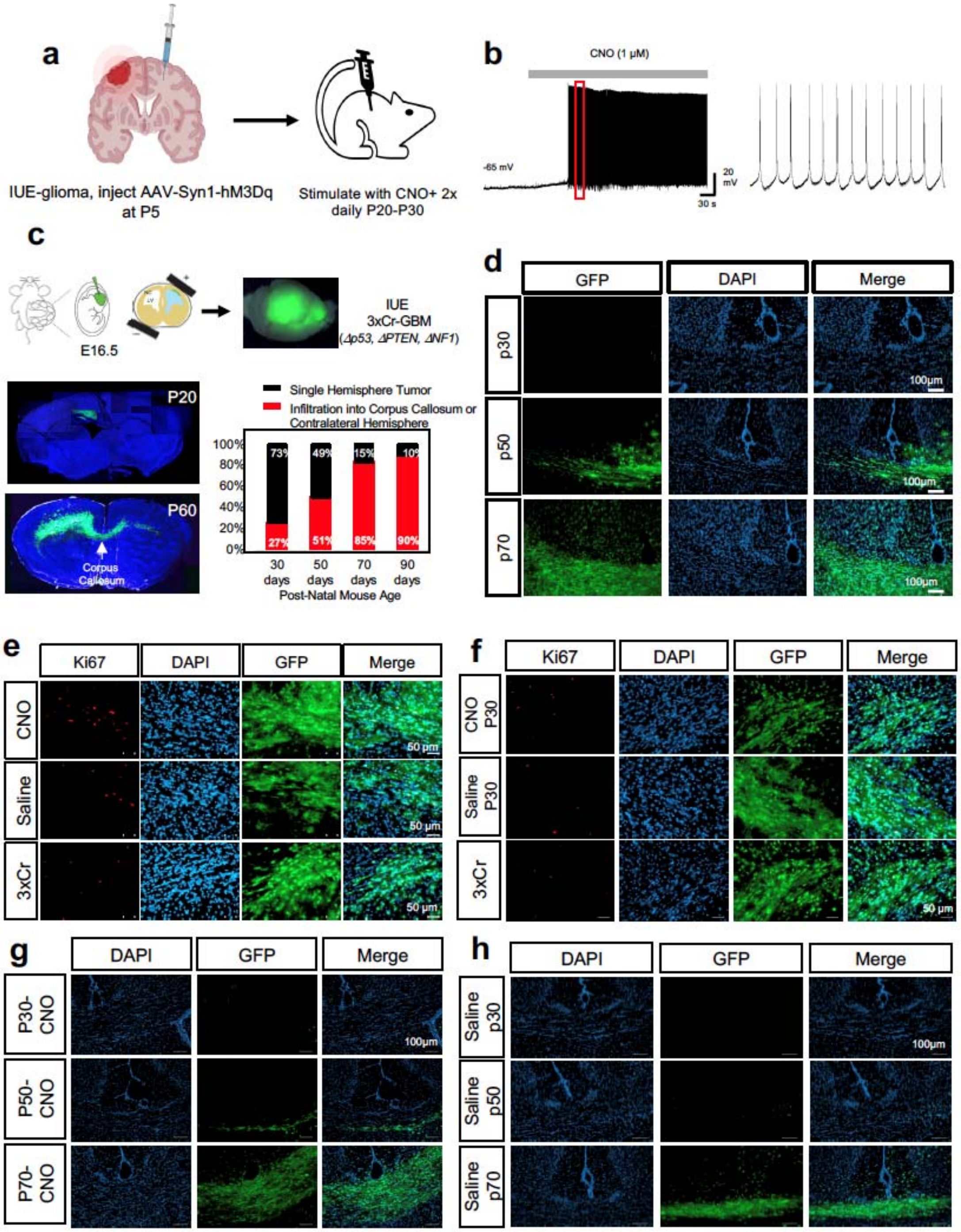


**Extended Data Figure 1. Contralateral cortical stimulation accelerates tumor infiltration**

**a.** Schematic depicting injection of AAV into the contralateral cortex. Brains were infected with AAV containing a pAAV-Syn1-hM3Dq-2a-mCherry cassette. Syn1 is a pan-neuronal promoter, thus all neurons infected expressed the hM3Dq-2a-mCherry cassette. **b.** Electrophysiology measuring neural activity in response to CNO treatment on mouse brain sections to confirm DREADD activity. **c.** Schematic of intra-uterine electroporation model of GBM. Cocktails include 3 guides targeting p53, PTEN, and NF1. Representative images of regular tumor brains from 20 day old mice versus 60 days old. Tumor brain slices were stained with Hoechst, and native GFP fluorescence was utilized to visualize tumor. Time course of contralateral cortical infiltration. Log-regression of 3xCr tumors demonstrated that infiltration correlates strongly with time (log-regression, Chi-square = 23.38, df = 1, p value <0.0001, $n_{30}$ =22, $n_{50}$=19,$n_{70}$=21, $n_{90}$=10) with half of cases appearing around 55-60 days. **d.** Representative images of infiltrating tumor across the midline at each respective timepoint. Frozen sections were stained with Hoechst and native GFP fluorescence was utilized to visualize tumor. **e.** Representative Ki67 staining of CNO versus Saline versus 3xCr IUE sections taken from 30 day old tumor brains. **f.** Representative Ki67 staining of CNO-only control, without hM3Dq, Saline, and 3xCr IUE sections taken from 30 day old tumor brains. **g-h.** Representative images of infiltrating tumor across the midline at each respective timepoint from the CNO-only control experiments (**g**) and saline only control (**h**).

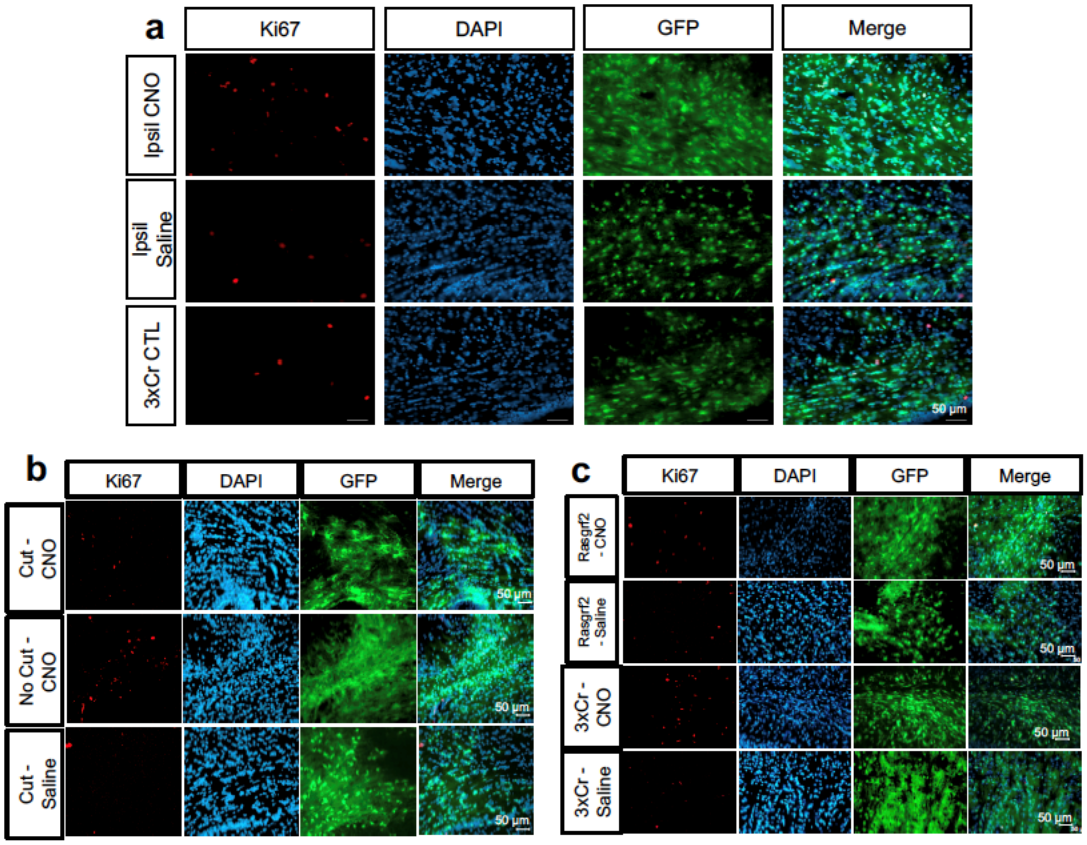


**Extended Data Figure 2. Ipsilateral stimulation does not promote early tumor infiltration**

**a.** Representative Ki67 staining of CNO versus Saline versus 3xCr IUE sections taken from 30 day old tumor brains after ipsilateral stimulation. **b.** Representative images of corpus callosum cut tumors stained for Ki67 (quantification in main figure). Tumor samples were prepared as described above, using native GFP to image tumor. **c.** Representative images of Rasgrf2-dCre tumors stained for Ki67 (quantification in main figure). Tumor samples were prepared as described above, using native GFP to image tumor. Levels of statistical significance are indicated as follows: * ($p < 0.05$), ** ($p < 0.01$), *** ($p < 0.001$), **** ($p < 0.0001$).

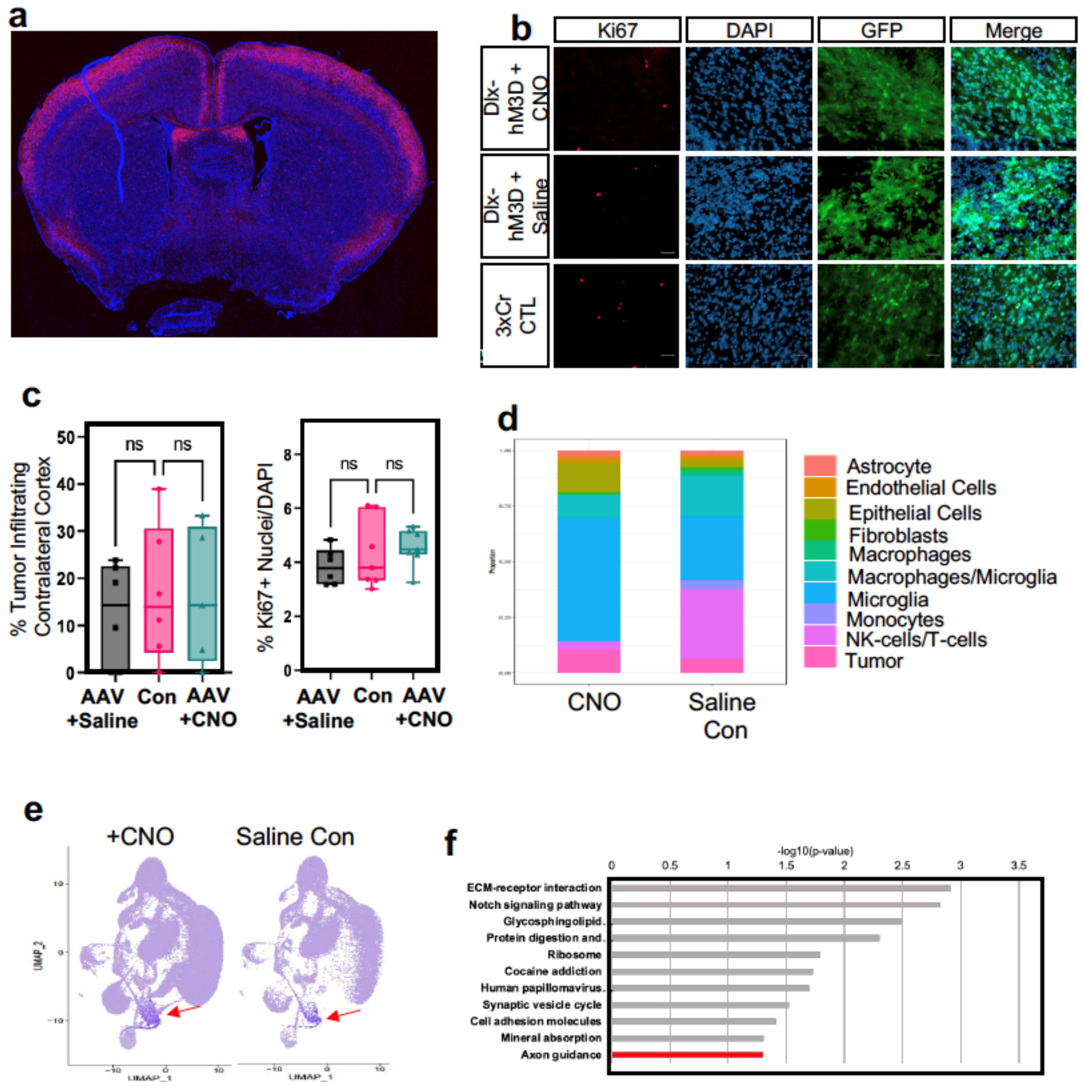


**Extended Data Figure 3. Representative Image of Rosa-LSL-TdTom + Rasgrf2-dCre**

**a.** Representative image of Rosa-LSL-TdTom + Rasgrf2-dCre mouse brain harvested 2 days post Trimethoprim treatment. Cells from layer 2/3 of the cortex show clear TdTom labelling. **b.** Quantification of tumor infiltration and Ki67 expression at P30; Infiltration was quantified based on the presence of tumor cells in contralateral cortex and analyzed via one-way ANOVA. Box and whisker plots outline minimum, maximum, and interquartile ranges. Data derived from 3xCr CTL (n=7), 3xCr + DLX-hM3Dq-mCh + Saline (n=6), 3xCr + DLX-hM3Dq-mCh + CNO (n=7) tumors with an average of 18 coronal slices analyzed per tumor. Percent coronal sections with infiltrating tumor was analyzed via one-way ANOVA and DLX-DREADD+CNO tumors showed no difference in proliferation to Saline treated (p-value = 0.1823) or control tumors (p-value = 0.5328) or infiltration at p30 to Saline (p-value = 0.8880) or control tumors (p-value = 0.9981). **c.** Representative images from IUE-HGG tumors at P30 demonstrating Ki67 expression after activation of inhibitory neurons with AAV-Dlx5/6-hM3Dq in the cortex contralateral to the tumor (CNO) and saline treated controls. **d.** Stacked bar chart of SingleR labeled populations percent representation in CNO and Saline single cell sequencing datasets. **e.** Feature plot of cluster of interest marker genes. Color represents a score assigned based on overall expression of marker genes. Seurat's AddModuleScore function was used to calculate these scores. Red Arrow denotes population of interest featured in Fig. 3B. **f.** GO terms of genes enriched in the spatial transcriptomics analysis of the leading edge of P50 mouse tumors (R6-R8).

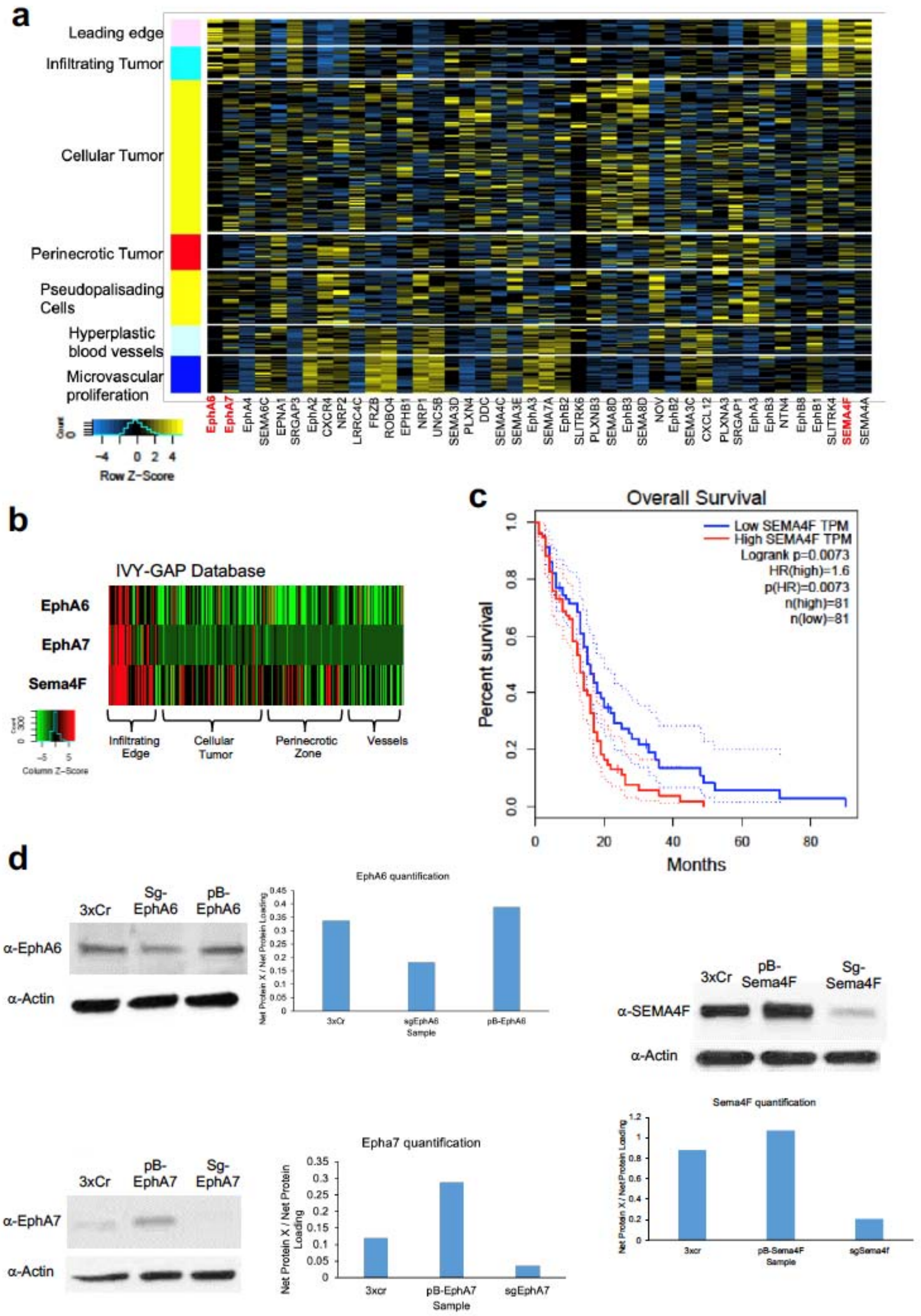


**Extended Data Figure 4. Expression characteristics of EphA6, EphA7, and Sema4F**

**a.** Heatmap of IVY-GAP expression data of all axon guidance genes included in the bar coded screen described in figure 4a. **b.** Heatmap of IVY-GAP (human) expression data for Sema4f, EphA6, EphA7. **c.** KM curve of GEPIA survival data. GBM patient cohort data were split based on high and low Sema4F expression and show a similar trend to our GOF and LOF in vivo data. **d.** Western blots of wild type, GOF, and LOF tumors. (pB-Sema4F is GOF; sgSema4F is LOF). Antibodies and concentrations used are outlined above. Western blots were quantified in ImageJ.

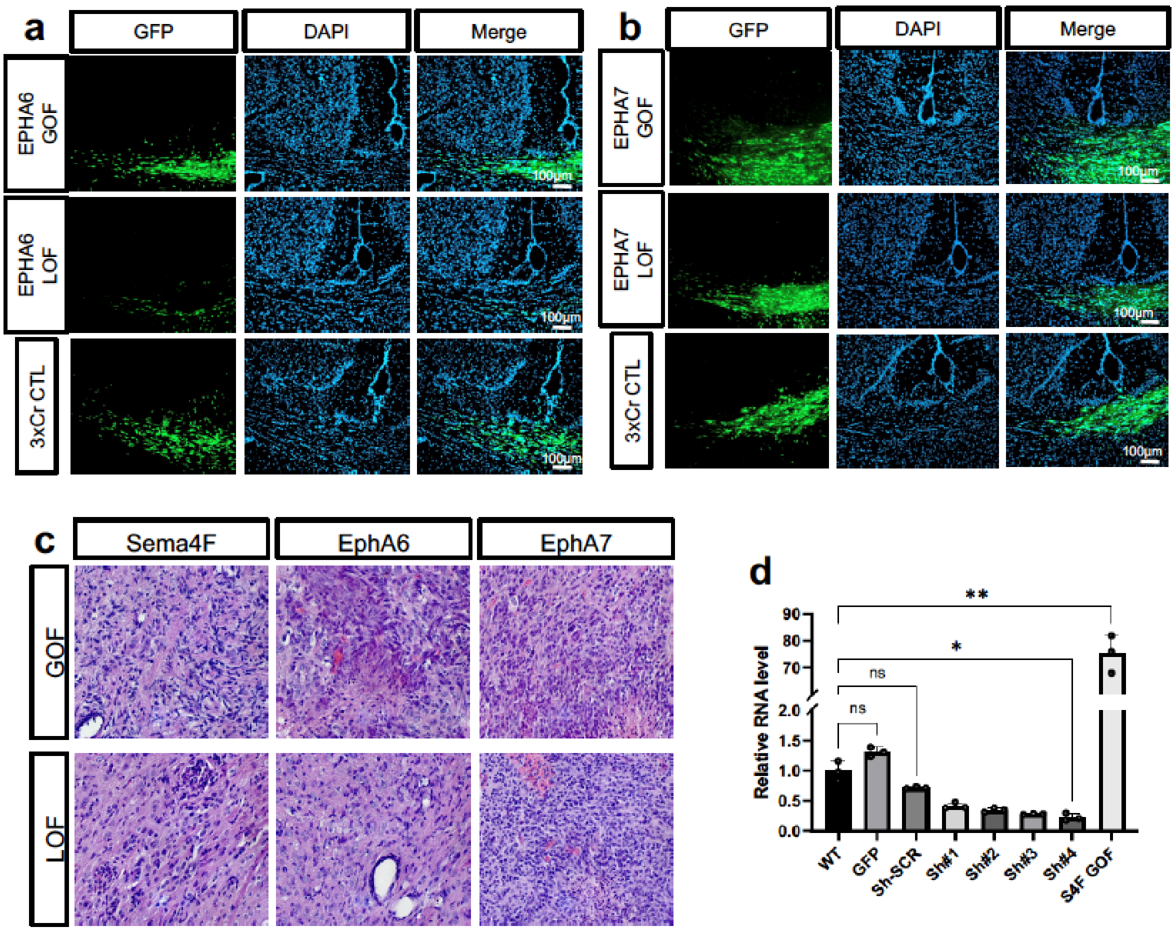


**Extended Data Figure 5. Infiltrating glioma from EphA6 and EphA7 GOF/LOF tumors**

**a.** Representative images of IUE-HGG GOF/LOF/CTL of EphA6 in 50 day old mice. Frozen sections were stained with Hoechst and native GFP was assessed for infiltration status into the contralateral cortex. **b.** Representative images of IUE-HGG GOF/LOF/CTL of Epha7 in 50 day old mice. Frozen sections were stained with Hoechst and native GFP was assessed for infiltration status into the contralateral cortex. **c.** Representative H&E staining of IUE-HGG tumors containing GOF and LOF of candidate genes. All tumors demonstrated high grade characteristics such as microvascular proliferation or necrosis regardless of GOF or LOF. **d.** qRT-PCR validation of Sema4F-GOF overexpression and shRNAi knockdown in human glioma cell lines. Values were plotted and analyzed by Brown-Forsythe and Welch analysis of variance (ANOVA) in Prism (v9.3.0). S4F-GOF showed significant upregulation compared to uninfected control (p-value = 0.0082) and GFP cells (p-value = 0.0083), while shS4F #4 showed significant downregulation compared to uninfected control (p-value = 0.0154) and shScramble cells (p-value = 0.0167).

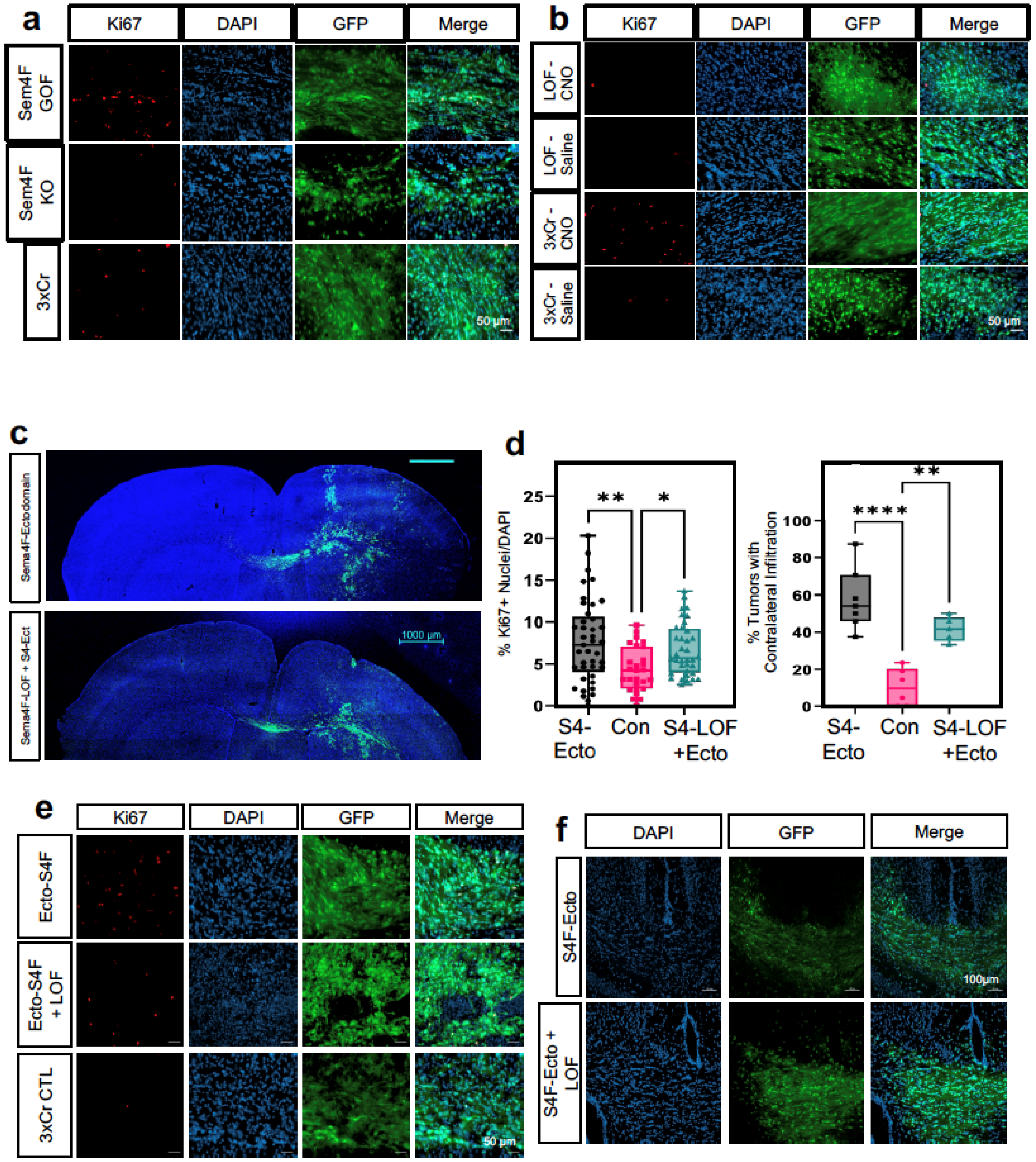


**Extended Data Figure 6. Representative Ki67 images from Sema4F manipulated tumors**

**a.** Representative images of Sema4F GOF/LOF and control tumors stained for Ki67 in 30 day old mice. Tumor samples were prepared as described above, using native GFP to image tumor. **b.** Representative images of Sema4F-LOF and control tumors treated with CNO or Saline and stained for Ki67 in 30 day old mice. Tumor samples were prepared as described above, using native GFP to image tumor. **c.** Representative images from IUE-HGG tumors at P30 demonstrating the extent of infiltration with Sema4F-ectodomain and Sema4F-ectodomain + Sema4F-LOF. **d.** Quantification of tumor infiltration and Ki67 expression at P30; Infiltration was quantified based on the presence of tumor cells in contralateral cortex and analyzed via one-way analysis of variance (ANOVA). Error bars represent standard error, data derived from N≥5 tumors derived from N≥5 mice, per condition, per timepoint. $^*P < 0.05$, $^{**}P < 0.01$, $^{***}P < 0.001$, $^{****}$ $P < 0.0001$, one-way analysis of variance (ANOVA) (**d**) **e.** Representative images of Sema4F-ectodomain and Sema4F-ectodomain + Sema4F-LOF and control tumors stained for Ki67 in 30 day old mice. **f.** Representative images of infiltrating tumor across the midline from Sema4F-ectodomain and Sema4F-ectodomain + Sema4F-LOF tumors in 30 day old mice.

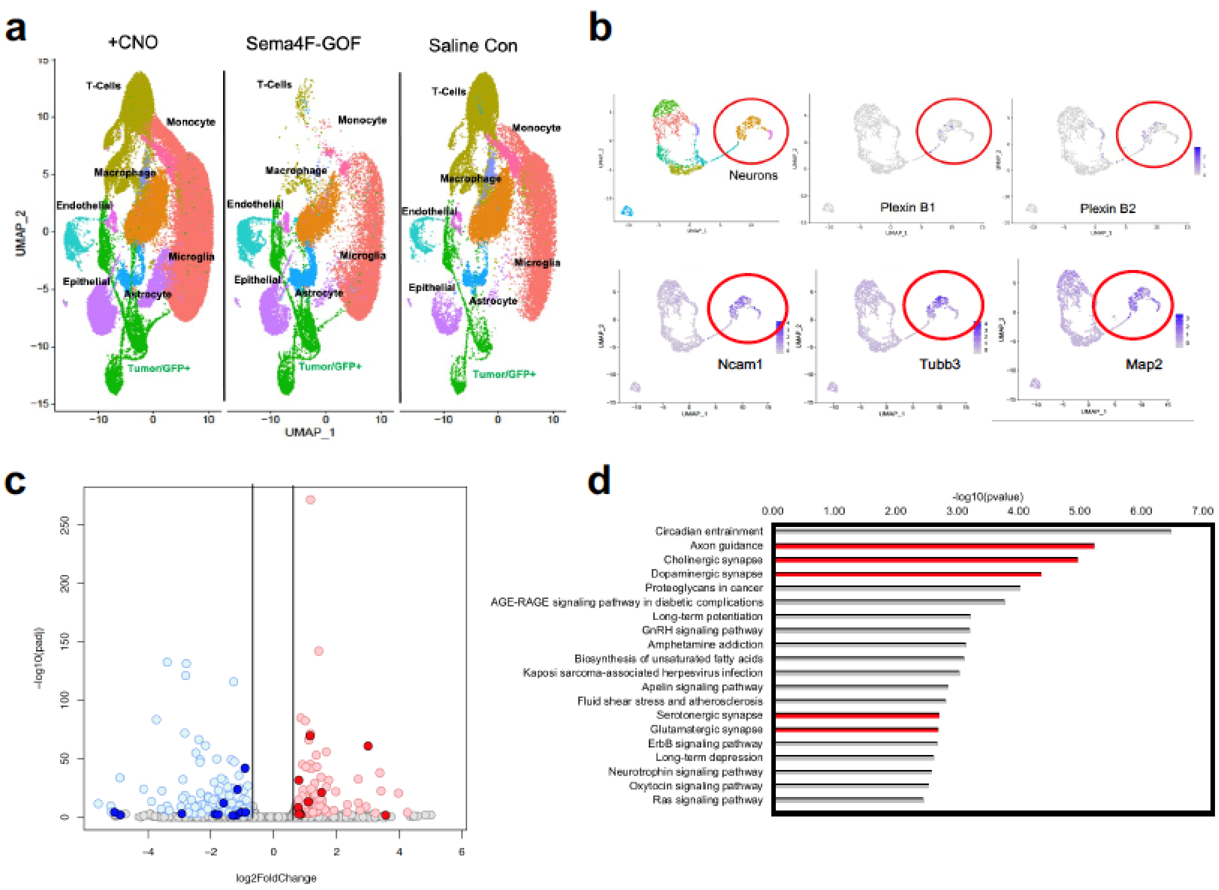


**Extended Data Figure 7. Single Cell RNA-Seq DimPlots of P50 IUE-HGG from CNO, Sema4F, and saline controls**

**a.** UMAP plot of full single cell datasets from CNO and Saline treated and Sema4F GOF tumors. Tumors were isolated and sequenced as outlined above. All clusters were represented in all datasets, though their relative abundance varied. **b.** Single cell RNA-Seq analysis of from IUE-glioma model, sub-clustering on non-tumor, GFP-negative cells. Red circle denotes neuronal populations, marked by Map2, Tubb3, and Ncam. Note that Plexin B1 and B2 expression is enriched in neuronal populations, denoted by purple dots. **c.** Volcano plot depicting the differentially expressed genes (DEGs) between human glioma cell lines overexpressing Sema4F (Sema4F-GOF) and controls. For downregulated genes with blue color: padj<.001 & log2FoldChange<(-0.75); for upregulated genes in red: padj<.001 & log2FoldChange>0.75) **d.** GO analysis of DEGs upregulated in the Sema4F-GOF glioma cell lines.

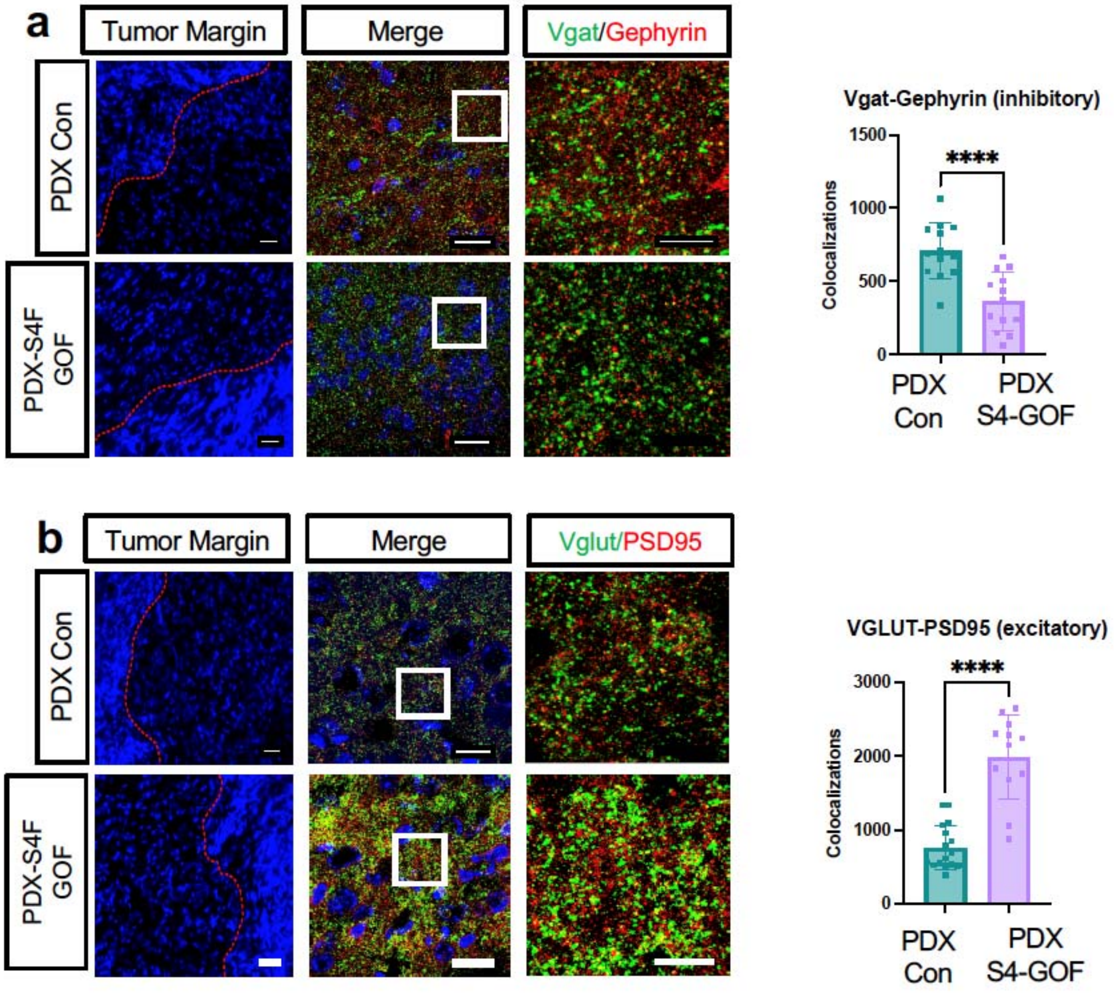


**Extended Data Figure 8. Staining for synaptic markers in mice bearing PDX-Sema4F-GOF tumors.**

**a.** Antibody staining of inhibitory synapses (VGAT-Gephryin) and **b**.excitatory (Vglut2-PSD95) synapse from P50 mouse brains at peritumoral margins from PDX tumors ovexpression Sema4F and control; box denotes zoomed in region in adjacent panel (20X, 63X, and 200X magnification left to right; scale bar left to right: 50 um, 20 um and 10 um). Quantification of synaptic staining derived from 3 separate tumors for each condition. Error bars represent standard error, data derived from 3 tumors derived from 3 mice with n≥15 fields analyzed and quantified, per condition. **** $P < 0.0001$, one-way analysis of variance (ANOVA)

**Extended Data Table 1**

GO-term analysis of upregulated genes in CNO treated mouse dataset versus Saline treated mouse dataset. Positive markers were found for CNO treated samples, and these were analyzed via Enrichr using the KEGG 2021 Human gene set.

**Extended Data Table 2**

Metadata for the final Seurat object containing CNO, Saline, and Sema4F-GOF scSeq

**Extended Data Table 3**

GO-term analysis of upregulated genes in subcluster of interest (Fig 3) in CNO treated mouse dataset versus Saline treated mouse dataset. Positive markers were found for CNO treated samples, and these were analyzed via Enrichr using the KEGG 2021 Human gene set.

**Extended Data Table 4**

GO-term analysis of upregulated found in the leading edge (R6-R8) of the spatial transcriptomics analysis from Figure 3. GO terms derived from Enrichr using the KEGG 2021 Human gene set

**Extended Data Table 5**

List of Axon Guidance genes used in the screen and their corresponding barcodes

**Extended Data Table 6**

GO-term analysis of upregulated genes in sub-subcluster of interest (Fig 5) in CNO treated and Sema4F GOF mouse dataset versus Saline treated mouse dataset. Positive markers were found for the cluster of interest, and these markers were analyzed via Enrichr using the KEGG 2021 Human gene set.

**Extended Data Table 7**

List of GO terms from human cell lines overexpressing Sema4F compared to controls. Analysis was performed with Enrichr using the KEGG 2021 Human gene set.

**Extended Data Table 8**

List of DEGs from human cell lines overexpressing Sema4F compared to controls. Analysis was performed with Enrichr using the KEGG 2021 Human gene set.
